# Endothelial cell-intrinsic NOD2 signaling regulates the intestinal immune response through the generation of effector and memory T cells

**DOI:** 10.1101/2025.09.06.674650

**Authors:** Boyan K. Tsankov, Madison Denney, Charles Carr, Jennifer SY. Ahn, Derek KL. Tsang, Giuliano Bayer, Lauren Baerg, Alexander Luchak, Nayanan N. Nathan, Tapas Mukherjee, Elaine Tam, Bana Samman, Jennifer Gommerman, Stephen E. Girardin, Dana J. Philpott

## Abstract

Crohn’s disease (CD) is marked by vascular endothelial dysfunction and aberrant T cell immunity in the gastrointestinal tract. However, mechanistic understanding is lacking of how CD-associated gene variants, particularly those that compromise NOD2 function, impact the intestinal vascular endothelium and orchestration of T cell immunity. Here, we find that NOD2, when triggered by its ligand, muramyl dipeptide, is unique among pattern recognition receptors in its ability to induce endothelial cell expression of chemokines and immune adhesion molecules. Consequently, NOD2 signaling promoted T cell homing specifically to gut-associated lymphoid tissue during intestinal infection. Endothelial cell-specific deletion of *Nod2* resulted in fewer effector and memory T cells in the small intestine, impacting the host’s ability to clear secondary infection. Together, our findings suggest that vascular endothelial cell expression of NOD2 coordinates intestinal immune responses, and that CD-associated loss of NOD2 function promotes aberrant inflammation due to alterations in the magnitude and specificity of host defense within the intestine.

## Introduction

Crohn’s disease (CD) is an inflammatory bowel disease (IBD) marked by severe inflammation of the gastrointestinal tract^1^. Although the etiology of CD remains incompletely understood, disease development is thought to involve a strong genetic predisposition, with polymorphisms in more than 150 genes resulting in increased risk of CD^2^. The largest genetic risk factor for CD development in Caucasian populations are mutations in nucleotide oligomerization-domain containing protein (*NOD2*) that result in a non-functional NOD2 protein^3^. *NOD2* encodes an intracellular pattern recognition receptor (PRR) that recognizes muramyl dipeptide (MDP) – a peptide fragment of bacterial peptidoglycan^4^. Individuals homozygous for *NOD2* genetic variants have a 20-40-fold increased risk of disease manifestation^5^. Interestingly, even in the absence of *NOD2* mutations, recent experiments have shown decreased NOD2 activation due to reduced levels of fecal MDP in patients with CD compared to healthy controls^6^. Mechanistically, NOD2 dysfunction results in aberrant intestinal adaptive immunity. Indeed, NOD2 is classically known to be expressed in high amounts in conventional dendritic cells (cDCs)^7^, and the recognition of MDP by NOD2 is well-established to potentiate adaptive immune B and T cell responses in secondary lymphoid organs (SLOs) and tissues^8^. Linking this fact to the notion that decreases in NOD2-mediated signalling predispose individuals to CD, suggests a potential link between mutations in *NOD2* and dampened adaptive immune responses in the intestine.

Aberrant intestinal T cell immunity is thought to be one of the main drivers of CD^9^. Historically, examinations of the mechanisms underlying intestinal T cell functionality and dysregulation have mainly focused on T cell interactions with other antigen-presenting cells (APCs) such as DCs^10^, and macrophages^11^. However, there are few investigations into the ways in which non-immune stromal cells such as fibroblasts, lymphatic endothelial cells, and blood endothelial cells affect intestinal T cell immunity. Interestingly, early studies following the characterization of NOD2 indicated protein and transcript presence in blood endothelial cells (BECs)^12^. However, there have been no further investigations into the effects of NOD2 on the endothelium and associated effects on intestinal T cell immunity. Interestingly, gut-vascular barrier dysfunctions such as aberrant angiogenesis^13^, innate immune sensing^14^, and endothelial permeability^15^ have been implicated in inflammatory bowel disease (IBD). Given the requirement for the vascular endothelium in mediating T cell trafficking to mesenteric lymph nodes for T cell priming^16^, and subsequently to the intestinal parenchyma for effector T cell functions^17^, an investigation into the ways in which NOD2 mediates these processes is needed for a better understanding of the etiology of CD.

During intestinal infection, naïve T cells home to the mesenteric lymph nodes (mLN) by following a chemotactic gradient involving C-C motif chemokine ligand 19 and 21 (CCL19 and CCL21)^18^, and extravasate from the blood into the inductive site via binding to central adhesion molecules (CAMs) present on blood endothelial cells (BECs)^19,20^. Once in the mLN, naïve T cells recognize the cognate antigen presented by professional APCs and rapidly proliferate more than 1000-fold via clonal expansion^21^. The expanded T cells subsequently enter the intestinal tissue bed, where they aid in the clearance of infection and, importantly, establish a memory T cell pool for rapid protection following subsequent antigen encounter^22^. While MDP recognition by APCs is known to promote the generation of effector and memory antigen-specific T cells following antigenic challenge^8^, the contribution of vascular endothelial sensing of MDP by NOD2 to these processes is unknown.

In this study, we investigated how NOD2 within the vascular endothelium contributes to the adaptive immune response to an intestinal infection, using lymphocytic choriomeningitis virus (LCMV) as a model of robust effector and memory T cell generation^23^. Using adoptive transfer experiments, MHC-tetramers, and single cell transcriptomics, we uncover a novel role for endothelial cell-intrinsic NOD2-signalling in mediating effector and memory T cell responses in the intestine, and for effective control of subsequent viral infection. We establish that NOD2 within the vasculature of the mesenteric lymph nodes (mLN) is critical for the homing and subsequent priming of naïve T cells during early infection. Mechanistically, we further show that NOD2-mediated sensing of MDP by the vascular endothelium uniquely induces a chemokine and adhesion-molecule program that is critical for T cell homing to gut-associated lymphoid tissue. This work identifies a novel role for NOD2 in regulating endothelial function, fine tuning intestinal T cell immunity to protect against infection and modulating inflammation.

## Results

### Endothelial NOD2 facilitates the generation of intestinal effector and memory T cells following infection

Given the importance of T cell immunity to intestinal homeostasis and CD manifestation, we first wanted to assess the effects of NOD2-deficiency on antigen-specific ileal T cells. We chose to focus our examination on the ileum as CD driven by NOD2 mutations primarily results in ileal pathology^24^. To appropriately examine T cell immunity in its effector and memory stages in an antigen-dependent way, we needed an infection model that would allow us to track immunodominant T cell clones throughout the T cell lifecycle; from priming in the mesenteric lymph nodes (mLNs) to effector and memory functions in the ileum. We decided to use a strain of lymphocytic choriomeningitis virus infection (LCMV-Armstrong) that causes acute infection, which has been recently shown to infect the small and large intestines, leading to the clonal expansion of LCMV-specific T cells in the mLNs^23,25^. We confirmed that acute LCMV infection of littermate WT and *Nod2*^−/−^ C57Bl/6 mice induced robust CD4^+^ and CD8^+^ T cell expansion in the small intestinal lamina propria (SILP) 8 days after infection (**Figures 1A and 1B**). Interestingly, we found that this effect was dependent on NOD2, as *Nod2*^−/−^ mice had significantly decreased numbers of lamina propria total TCRβ^+^ and CD8^+^ T cells (**Figure 1B**). Furthermore, this phenotype extended to LCMV-specific CD4^+^ and CD8^+^ T cells at this timepoint, with *Nod2*^−/−^ mice exhibiting significantly fewer numbers of antigen-specific gp66-77 (CD4^+^) and gp33-41 (CD8^+^) tetramer-labelled cells (**Figure 1C**). No genotypic differences in intraepithelial lymphocyte (IEL) and splenic T cell responses were observed (**Figures S1A and S1B**), indicating a specialized role for NOD2 in lamina propria effector T cell generation. Transcriptional assessment of the intestinal chemotactic chemokine *Ccl25*^26^, and T cell maintenance cytokines *Il7, Il15*, and *Tgfb*^27^ did not reveal a downregulation of these factors in *Nod2*^−/−^ mice (**Figure S1C**), indicating that the observed decrease of SILP-tropic T cells in *Nod2*-deficient mice is likely not due to differences in local microenvironmental cues.

**Figure 1.**
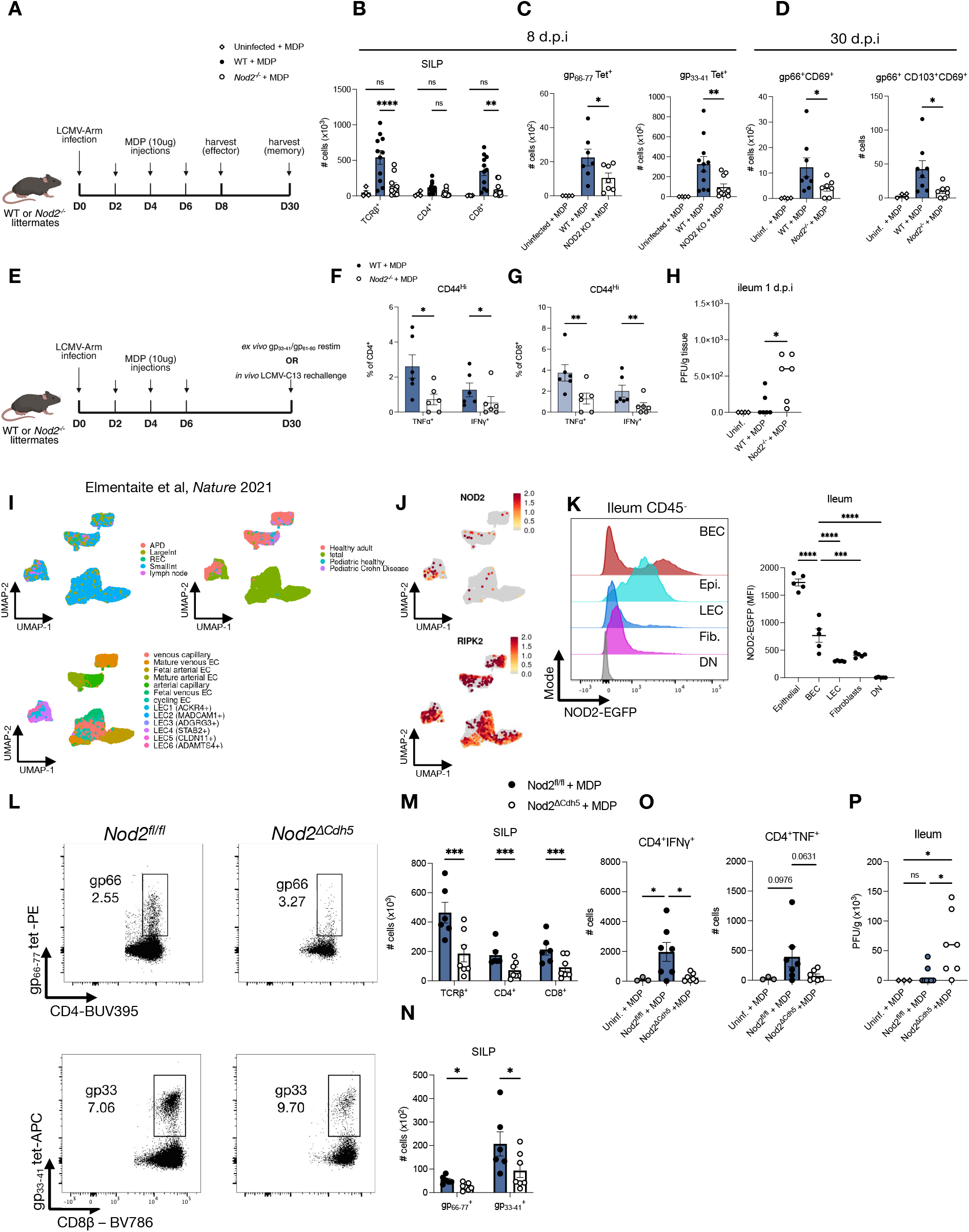
Endothelial NOD2 facilitates the generation of antigen-specific effector and memory T cells following acute intestinal infection. (A) Experimental schematic for the flow cytometry analysis of the number of effector and memory T cells in the SILP after LCMV-Armstrong infection. (B) Flow cytometry analysis of the number of total TCRβ^+^, CD4^+^, and CD8^+^ T cells 8 days following LCMV infection. n >= 4 mice per group; ordinary two-way ANOVA. (C) Flow cytometry analysis of the number of LCMV-specific CD4^+^ (left) and CD8^+^ (right) T cells in the SILP 8 days post LCMV-Armstrong infection. n >= 4 mice per group; ordinary two-way ANOVA. (D) Flow cytometry analysis of the number of LCMV-specific CD4^+^ T_RM_ cells in the SILP 30 days post infection. n >= 4 mice per group; ordinary one-way ANOVA. (E) Experimental schematic of the *in vitro* and *in vivo* LCMV rechallenge experiments. (F-G) Flow cytometric analysis of the *ex vivo* peptide recall potential of CD4^+^ (F) and CD8^+^ (G) memory T cells isolated from the SILP 30 days following LCMV-Armstrong infection of WT and *Nod2*^−/−^ littermates. N = 6 mice per group; ordinary two-way ANOVA. (H) Plaque assay of LCMV-C13 PFUs isolated from the terminal ileum of mice 24h post LCMV-C13 rechallenge *in vivo*. n >= 4 mice per group; ordinary one-way ANOVA. (I) Aggregate scRNA-seq UMAP plot of fetal, pediatric CD, pediatric healthy, and adult endothelial cells from appendix (APD), large intestine (LargeInt), rectum (REC), small intestine (SmallInt) and lymph nodes (lymph node) from gutcellatlas.org^28^. (J) FeaturePlot of *NOD2 and RIPK2* feature expression across endothelial cells. (K) Flow cytometric analysis of NOD2-EGFP expression in murine blood endothelial cells (BEC) epithelial cells (Epi.), lymphatic endothelial cells (LEC), fibroblasts (Fib.) and double negative (DN) cells within naïve ileal tissue. n = 5 mice; ordinary one-way ANOVA. (L) Representative flow cytometry plots of the number of LCMV-specific CD4^+^ and CD8^+^ T cells 30 days post infection of *Nod2*^*fl/fl*^ and *Nod2*^*ΔCdh5*^ littermates. (M-N) Flow cytometry analysis of the number of T cells (M) and antigen-specific T cells (N) of WT and *Nod2*^*ΔCdh5*^ littermates 30 days following LCMV-Armstrong infection. n = 6 mice per group; ordinary two-way ANOVA. (O) Flow cytometry analysis of the number of ileal IFNγ and TNF-producing CD4^+^ T cells 24 hours following rechallenge with LCMV-C13. Mice were infected with LCMV-Armstrong 30 days prior. No *ex vivo* restimulation or Golgi-transport inhibitors were added to capture cells uniquely activated by the secondary infection *in vivo*. n=3-7 mice per group; ordinary one-way ANOVA. (P) Plaque assay of LCMV-C13 PFUs isolated from the terminal ileum of mice 24h post rechallenge *in vivo*. n = 3-7 mice per group; ordinary one-way ANOVA. *p < 0.05; **p < 0.01; ***p < 0.001; ****p < 0.0001. Mean ± SEM depicted.

We next assessed the development of LCMV-specific memory T cells in the SILP 30 days after infection by MHC tetramer staining. Numbers of LCMV-specific CD4^+^ resident memory T cells, characterized by their expression of CD69 and CD103, were increased by MDP treatment in NOD2 sufficient mice compared to *Nod2*^−/−^ animals (**Figure 1D**). Furthermore, *ex vivo* cognate peptide restimulation experiments 30 days post LCMV-Arm infection showed that intestinal memory CD4^+^ and CD8^+^ T cells from *Nod2*^−/−^ mice had decreased ability to respond to secondary peptide rechallenge specifically in the small intestine (**Figures 1F and 1G**), but not in the colon or spleen (**Figures S1D and S1E**). These findings indicate that *Nod2*^−/−^ mice likely have a decreased ability to clear secondary small intestinal infections in a T cell-dependent manner. To provide concrete evidence for this, WT and *Nod2*^−/−^ mice were re-infected with LCMV-Clone 13 (LCMV-C13; a heterotype of LCMV-Arm) 30 days post LCMV-Arm infection and assessed the viral titres in the small intestine (**Figure 1E and 1H**). *Nod2*^−/−^ animals had greater viral titres in the small intestine relative to WT control mice (**Figure 1H**), indicating a role for NOD2 in the control of secondary infections.

We were particularly interested in examining the role of the vascular endothelium to these phenotypes, as the contribution of non-immune stromal cells to CD etiology remains relatively understudied. Furthermore, given the necessity of the vascular endothelium to T cell accumulation within tissue, we examined whether NOD2 is expressed within human endothelial cells, by making use of publicly available single-cell RNA sequencing (scRNA-seq) datasets^28^. Interestingly, NOD2, and its downstream adaptor protein RIPK2 were expressed in BECs and lymphatic endothelial cells (LECs) of adult, fetal, and pediatric intestinal tissue (**Figures 1I and 1J**). We wanted to verify these results by making use of *Nod2*^+/−^ mice in which one allele of *Nod2* has been disrupted with an EGFP cassette^29^, and probing the expression of NOD2 among epithelial cells as a positive control, and non-immune stromal cells in the ileum by flow cytometry. Our results indicated that BECs robustly express NOD2 to significantly greater extents than other stromal cells in the intestine (**Figure 1K**). To confirm NOD2’s role in endothelial cells (ECs) *in vivo*, we leveraged Cdh5^CRE^ mice^30^ to specifically delete *Nod2* in endothelial cells by crossing these animals with *Nod2*^*fl/fl*^ mice (*Nod2*^*ΔCdh5*^).We next examined the generation of memory T cells following LCMV infection in these mice. *Nod2*^*ΔCdh5*^ mice showed significantly decreased numbers of antigen-specific CD4^+^ and CD8^+^ T cells in the small intestinal lamina propria 30 days post-acute LCMV infection (**Figures 1L-N**). Furthermore, these cells were confirmed to have a memory phenotype indicated by their expression of the memory markers CD69 and CD44 (**Figure S1F and S1G**). We next examined the recall potential of LCMV-specific memory CD4^+^ T cells generated in *Nod2*^*ΔCdh5*^ ilea *in vivo. Nod2*^*fl/fl*^ and *Nod2*^*ΔCdh5*^ mice were infected with LCMV-Armstrong, and, 30 days later, were re-infected with LCMV-C13 for 24 hours. Our results show that *Nod2*^*ΔCdh5*^ mice produce fewer pro-inflammatory Th1-like CD4^+^ T cells following secondary viral rechallenge than control littermates (**Figure 1O**). Furthermore, EC-specific deletion of *Nod2* significantly reduced the clearance of ileal LCMV-C13 following rechallenge (**Figure 1P**). Together, these data indicate that EC-intrinsic NOD2 signalling is required for effector and memory T cell generation in the small intestinal lamina propria, and for protection during secondary pathogenic intestinal infections.

### NOD2 engagement is sufficient for GALT adhesion molecule expression and antigen-specific T cell generation in microbiota-depleted mice

Given that NOD2 is an intracellular PRR of bacterial MDP, we next wanted to assess if the microbiota is required for induction of LCMV-specific T cell accumulation in the ileum. Furthermore, we wanted to assess whether NOD2 engagement via MDP administration is sufficient for intestinal antiviral T cell immunity. To do this, naïve WT mice were given oral doses of a cocktail of broad-spectrum antibiotics (ABX) in their drinking water for 3 weeks to deplete the intestinal microbiota, and subsequently infected with LCMV-Armstrong. We further evaluated whether supplemental MDP, delivered by intraperitoneal injections, could rescue NOD2-regulated immune responses in ABX-treated mice (**Figures 2A and 2B**). ABX-mediated depletion of the microbiota significantly reduced the numbers of total, and LCMV-specific CD4^+^ and CD8^+^ T cells (**Figures 2C-G**) in the small intestinal lamina propria (SILP) 7 days after infection. Interestingly, similar effects were observed in the spleen, implicating a far-reaching effect of the microbiota in antiviral immunity beyond the intestine (**Figure 2C-G**). Strikingly however, MDP administration rescued total CD8^+^ T cell, and LCMV-specific CD4^+^ and CD8^+^ T cell numbers of ABX-treated mice specifically in the SILP but not in the spleen (**Figure 2C-G**) implicating NOD2 as an indispensable regulator of antiviral T cell immunity specifically in GALT. Since our previous data implicated EC-intrinsic NOD2 in ileal T cell accumulation following infection (**Figures 1L-N**), we reasoned that microbiota-regulated differences in adhesion molecules may drive some of those effects. Indeed, the presence of microbiota was necessary for maintenance of *Vcam1* and *Madcam1* in the ileum following LCMV infection (**Figures 2H and 2I**). Strikingly, MDP was sufficient to rescue the ABX-mediated downregulation of adhesion molecule transcripts in the small intestine (**Figures 2H and 2I**). Given the importance of MAdCAM-1 for T cell homing specifically to mucosal sites such as the mLN, Peyer’s patches, and intestinal parenchyma^31^, we further verified if microbiota is required for MAdCAM-1 expression in the mLN, and whether MDP is sufficient for its protein expression in this compartment. Microscopic assessment of the mLN endothelium revealed that the microbiota is necessary for MAdCAM-1 expression in this compartment, with ABX-treated mice displaying significantly reduced protein colocalization with CD31, a marker of endothelial cells^32^, relative to non-ABX treated, control mice (**Figures 2J and 2K**). Strikingly, NOD2 stimulation via MDP was sufficient to rescue homeostatic MAdCAM-1 expression in the mLN endothelium (**Figures 2J and 2K**). Taken together, these results establish that the microbiota-derived MDP-NOD2 signalling axis is sufficient to induce optimal GALT T cell responses against viral infection, likely via enhancing endothelial functions related to immune cell homing.

**Figure 2.**
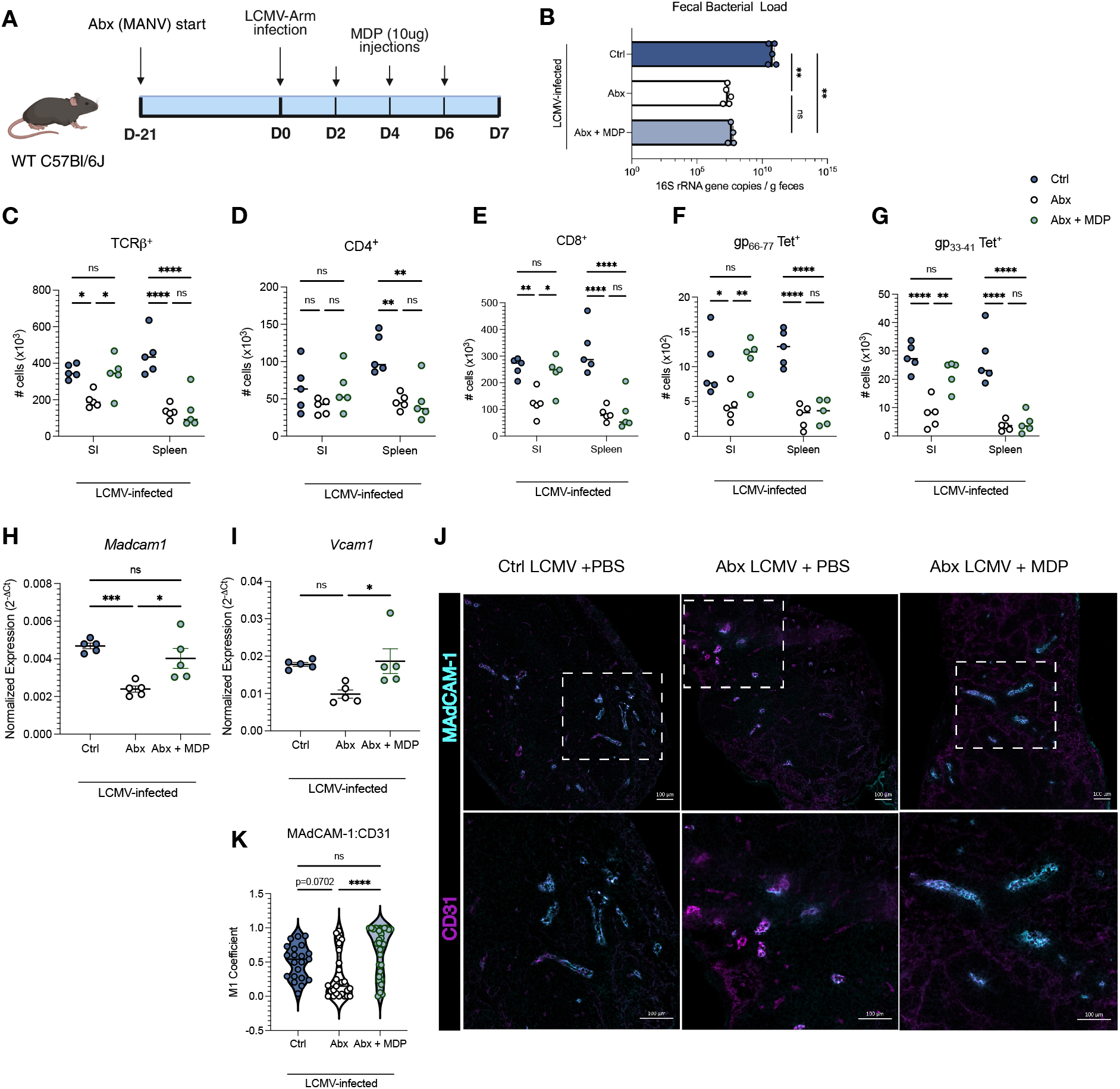
NOD2 engagement is sufficient for GALT adhesion molecule expression and antigen-specific T cell generation in microbiota-depleted mice. (A) Experimental schematic of the experimental design and administration of the metronidazole, ampicillin, neomycin, and vancomycin antibiotic cocktail. (B) qRT-PCR analysis of the 16S rRNA gene copies of ABX-treated mice at the experimental endpoint. (C-E) Flow cytometry population analysis of bulk TCRβ^+^, CD4^+^, and CD8^+^ T cells in the SILP and spleen of LCMV infected mice. n = 5-6 mice per group; ordinary one-way ANOVA. (F-G) Flow cytometry population analysis of LCMV-specific CD4^+^ gp_66-77_ tetramer^+^ (F), and CD8^+^ gp_33-41_ tetramer^+^ (G) T cells in the SILP and spleen of LCMV infected mice. n = 5-6 mice per group; ordinary one-way ANOVA. (H-I) qRT-PCR analysis of *Madcam1*(H) and *Vcam1* (I) expression in whole small intestinal tissue. n = 5-6 mice; ordinary one-way ANOVA. (J-K) Confocal microscopy and associated colocalization coefficients of CD31 and MAdCAM-1 in infected mouse mLNs. Each point represents one CD31^+^ from 5-6 mice per group; ordinary one-way ANOVA. *p < 0.05; **p < 0.01; ***p < 0.001; ****p < 0.0001. Mean ± SEM depicted.

### NOD2 is expressed in blood endothelial cells of the mesenteric lymph nodes, and its engagement induces a leukocyte recruitment program

We were interested in examining how NOD2 affects BECs within the mLN, given the fact that the mLN is the most upstream site of an intestinal T cell response, and also our observation that NOD2 stimulation induces mLN BEC MAdCAM-1 during infection (**Figures 2J and 2K**). To verify whether NOD2 is likely to have BEC-intrinsic functions in the mLN, we first asked whether mLN BECs express NOD2 by flow cytometry. Strikingly, we found the highest NOD2 expression within mLN BECs, which expressed even higher levels of NOD2 than conventional dendritic cells (cDCs – reported to express high amounts of NOD2^33^) (**Figure S2A**). We found that this high expression of NOD2 among BECs was not restricted to the mLN, as cervical, peripheral, and inguinal LN BECs expressed similarly high levels of NOD2 (**Figure S2B**). Despite this, however, we found regional variation in the stromal cell types that preferentially express NOD2 across different SLOs. NOD2 was preferentially expressed in the BECs of GALT SLOs (mLN and Peyer’s patches), and in the fibroblastic reticular cells (FRCs) of pLNs, iLNs, and cLNs (**Figure S2C-E**), suggesting potential differences in stromal-intrinsic NOD2-mediated effects between GALT and non-GALT LNs. Altogether, these data identify that BECs in GALT SLOs highly express NOD2 and, thus, may contribute to coordinating NOD2-dependent T cell responses in the mLN.

To probe mechanisms specifically within the mLN stroma by which NOD2 engagement may mediate LCMV-specific T cell responses, we performed scRNA-seq on stromal-enriched mLN fractions of untreated (NT) and LCMV-infected WT and *Nod2*^−/−^ mice treated 6 days prior with MDP (**Figures 3A and S2F**). ScRNA-seq corroborated our previous observation that *Nod2* is most highly expressed within BECs (**Figures 3B and S2G**). However, as we did not observe any effects of NOD2-deficiency on the gross cellular composition of the infected mLN stroma (**Figures S3A and S3B**), further analyses focused on the transcriptional differences within BECs between WT and *Nod2*^−/−^ mice. To identify transcriptional programs that are induced during inflammation within BECs, we performed differential gene expression analyses between BECs from untreated versus LCMV-infected mLNs. Expectedly, infection induced key chemokine expression, such as *Cxcl9, Cxcl10, Ccl2*, and *Ccl19* within BECs, and gene-set enrichment analysis (GSEA) against Gene Ontology Biological Processes revealed that infection augments gene signatures of leukocyte migration and cell-cell adhesion – processes that are critical for T cell infiltration into SLOs during inflammation (**Figures 3C and 3D**). Interestingly, differential gene expression (DGE) analyses of infected WT and *Nod2*^−/−^ BECs showed decreased chemokine transcription in infected *Nod2*^−/−^ BECs relative to WT BECs indicating that endothelial-specific NOD2 engagement is likely important for inducing a T cell chemoattractant program (**Figures 3E and 3F**). These results were corroborated by GSEA that indicated that WT BECs have an enrichment of transcriptional programs relating to taxis and regulation of cell migration during infection (**Figure 3G**). Importantly, this NOD2-mediated enhancement of a chemotactic gene signature was restricted to BECs, with FRCs and LECs displaying minimal changes in gene expression between infected WT and *Nod2*^−/−^ mice (**Figure S3C and S3D**). Together, these results indicate that NOD2 engagement drives a transcriptional phenotype in BECs conducive to T cell recruitment to SLOs.

**Figure 3.**
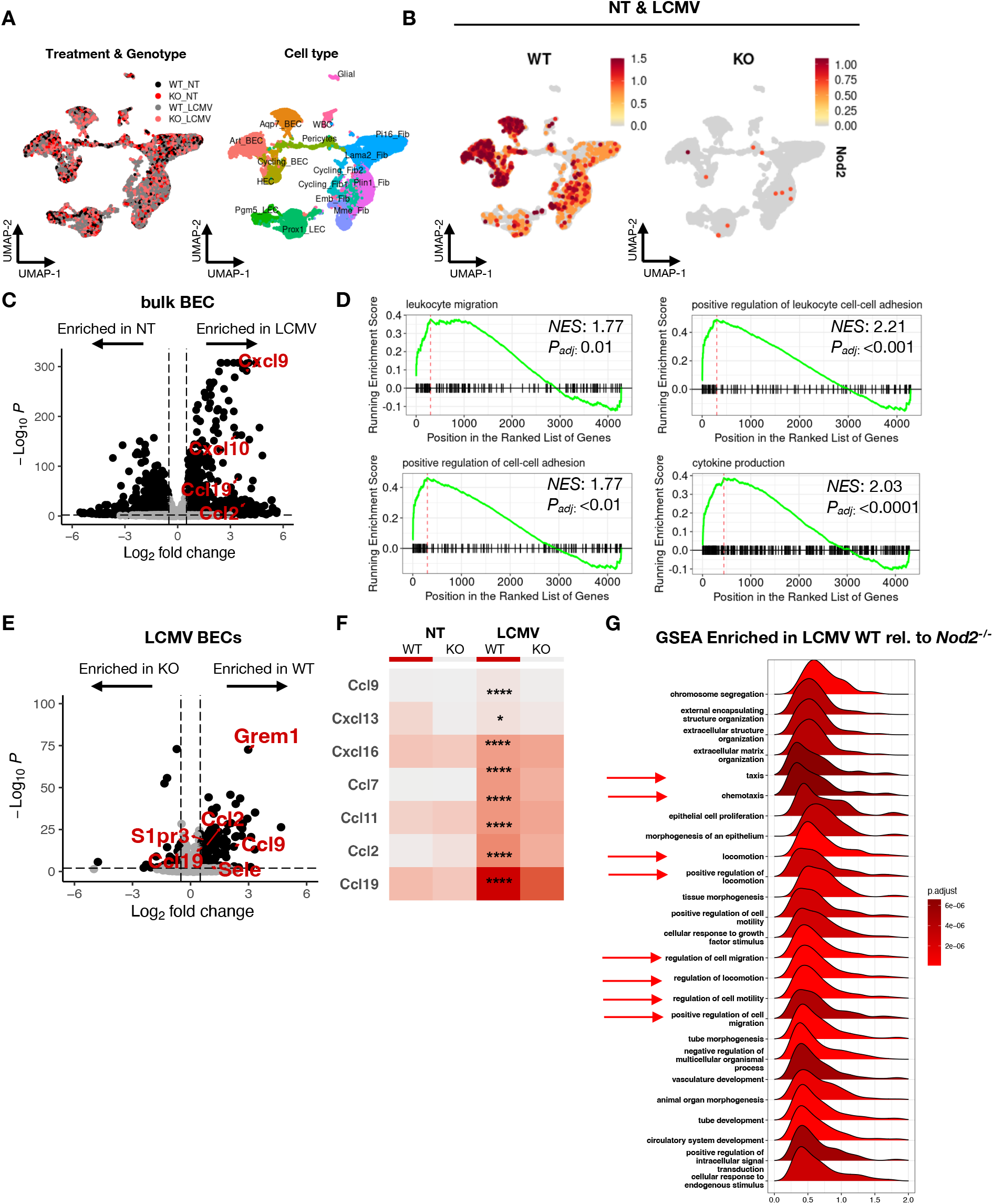
NOD2 is highly expressed in mLN blood endothelial cells and its engagement induces a leukocyte recruitment program. (A) Aggregate scRNA-seq UMAP plot of mLN stromal cells from naïve (NT) and LCMV-infected (6 days) WT and *Nod2*^−/−^ littermates. (B) FeaturePlot of *Nod2* feature expression across stromal cells of NT and LCMV-infected mice. (C) Volcano plot of DEGs between mLN BECs of NT and LCMV-infected mice. BECs were subsetted from the WT mLN stromal cell dataset. (D) Gene set enrichment plots of leukocyte migration, positive regulation of leukocyte cell-cell adhesion, positive regulation of cell-cell adhesion, and cytokine production biological process terms upregulated in infected BECs relative to NT BECs (E) Volcano plot of DEGs between infected WT and *Nod2*^−/−^ mLN BECs. (F) Heatmap of significantly enriched chemokines in BECs of infected WT mice relative to infected *Nod2*^−/−^ mice. Stars of significance symbolize p-values of significant differences between infected WT and infected *Nod2*^−/−^ BECs. (G) Gene-set enrichment analysis of the top 25 enriched gene set signatures in BECs of infected WT mice relative to infected *Nod2*^−/−^ mice. Gene gets related to leukocyte chemotaxis are highlighted with red arrows. *p < 0.05; **p < 0.01; ***p < 0.001; ****p < 0.0001. scRNA-seq data are pooled from n = 8 WT mice and n = 8 *Nod2*^−/−^ mice.

### Systemic administration of MDP leads to T cell accumulation in the mLN at homeostasis and during LCMV infection

Given the high *Nod2* expression pattern of mLN BECs and the induction of a transcriptional program of leukocyte recruitment following MDP administration, we next wanted to ask if NOD2 engagement within the mLN induces increased T cell accumulation at homeostasis and during early LCMV infection. To assess the effect of NOD2 at homeostasis, we engaged NOD2 in WT mice via i.p injection of MDP, or PBS control (**Figures 4A-E**). We found a NOD2-dependent increase in both the cellularity (**Figure 4A**) and numbers of B as well as CD4^+^ and CD8^+^ T cells in the mLN (**Figures 4B-E**). Interestingly, despite a significant increase in the total cellularity of the iLN (**Figure S4A**), we observed modest increases in the amounts of B and T lymphocytes following MDP administration (**Figure S4B**). Similarly, we did not observe any effect of NOD2 engagement on the splenic cellularity (**Figures S4C and S4D**), again suggesting a specialized role for NOD2 within the mLN to drive lymphocyte accumulation within this SLO. Given that the mLN is the main site of intestinal T cell priming^17^, we next assessed the relative amounts of T cells in the small intestinal lamina propria (SILP) following treatment with MDP. We did not observe any differences in the amount of total and CD44^+^CD69^+^ resident-memory T cells (TRM) between wildtype (WT) and *Nod2*^−/−^ littermate animals (**Figure S4E**), supporting the idea that NOD2 engagement specifically drives T cell accumulation to the mLN.

**Figure 4.**
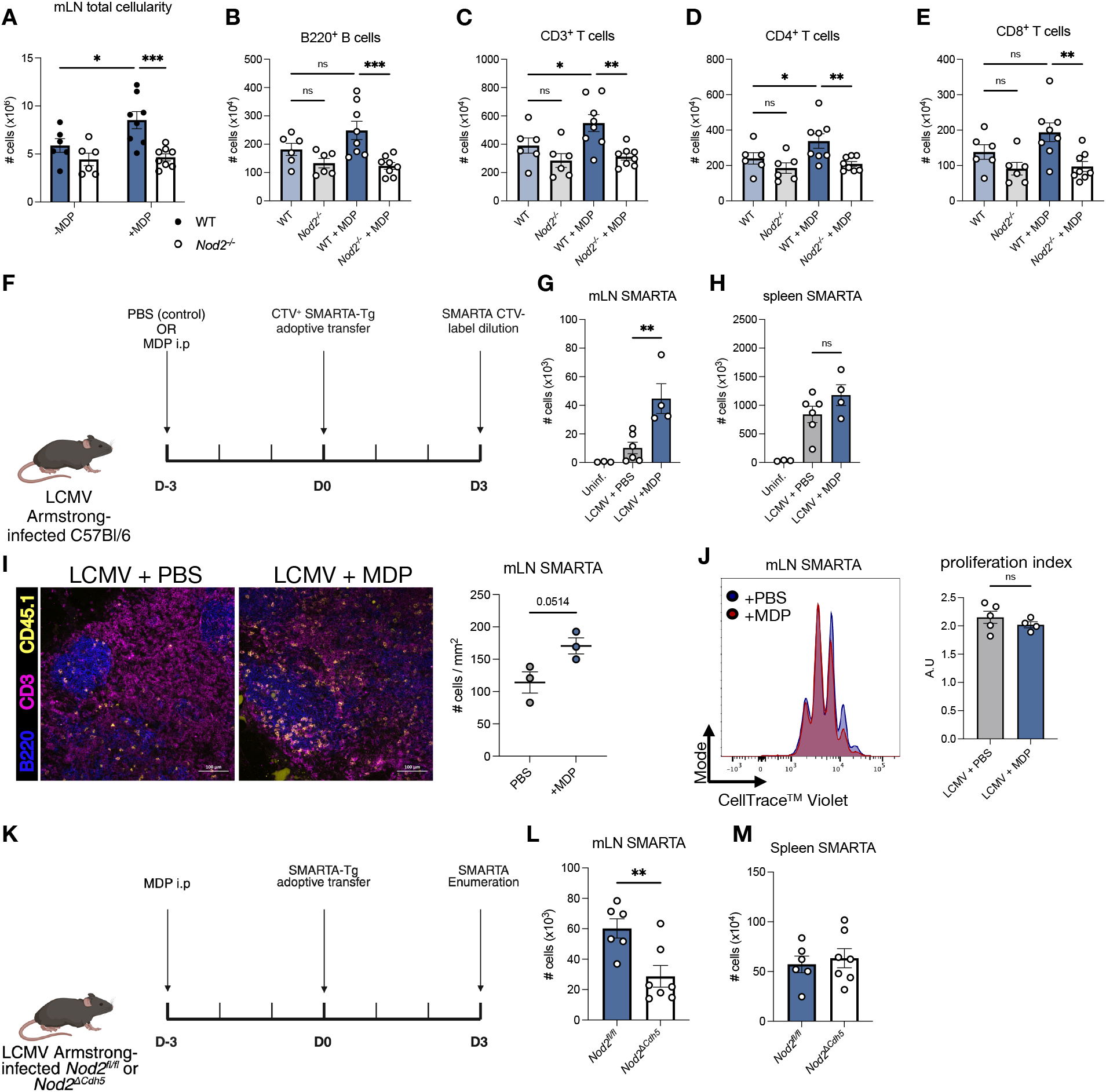
NOD2 engagement causes increased T cell accumulation in the mLN at homeostasis and during systemic LCMV infection. (A) Total live leukocyte cellularity of the mLN of WT and *Nod2*^−/−^ littermates. n > 5 mice per group; two-way ANOVA. (B-C) Flow cytometry analysis of the number of B (B) and T cells (C) in the mLN of WT and *Nod2*^−/−^ littermates. n > 5 mice per group; ordinary one-way ANOVA. (D-E) Flow cytometry analysis of the number of CD4^+^ T cells (D) and CD8^+^ T cells (E) in the mLN of WT and *Nod2*^−/−^ littermates. n > 5 mice per group; ordinary one-way ANOVA. (F) Experimental schematic of LCMV ± MDP infection of WT mice. Related to panels (G-J). (G-H) Flow cytometric analysis of the number of CD45.1 SMARTA-Tg CD4^+^ cells recovered from the mLN and spleen (H) of recipient mice 3 days post adoptive transfer. n >= 4 mice per group; ordinary one-way ANOVA. (I) Confocal microscopy and quantification of mLNs of LCMV-infected mice 3 days following adoptive transfer of CD45.1 SMARTA-Tg cells. Scale bar = 100µm. n = 3 mice per group; unpaired student’s t-test. (J) CellTrace™ violet dye dilution of mLN CD45.1 SMARTA-Tg CD4^+^ T cells and associated proliferation index 3 days following adoptive transfer. Proliferation index calculated using FlowJo. (K) Experimental schematic of SMARTA-Tg T cell accumulation experiments in WT and *Nod2*^*ΔCdh5*^ littermate mice. (L-M) Flow cytometric analysis of the number of adoptively transferred CD4^+^ SMARTA-Tg cells in the mLNs (L) and spleens (M) of recipient littermates. n = 6-7 mice per group; unpaired student’s t-test. *p < 0.05; **p < 0.01; ***p < 0.001; ****p < 0.0001. Mean ± SEM depicted.

We reasoned that the increased T cell accumulation in the mLN of WT animals following MDP priming could be due to several factors outside of homing, such as increased T cell activation, proliferation, or decreased cell death. To address the effects of NOD2 on these processes, we infected WT mice with LCMV-Arm with or without a NOD2 stimulus in the form of MDP (**Figure 4F**). Three days later, we adoptively transferred 5 x 10^5^ congenically-marked (CD45.1) naïve SMARTA-Transgenic CD4^+^ T cells (SMARTA) that have a T cell receptor (TCR) specific for the gp61-80 LCMV epitope. The transferred cells were also labelled with a CellTrace™ dye that enabled quantitative measurements of their proliferation by dye dilution. As expected, NOD2 engagement led to increased numbers of endogenous and SMARTA CD4^+^ T cells in the mLN, but not in the spleen (**Figures 4G-I**). Furthermore, with or without NOD2 stimulation, the adoptively transferred naive SMARTA-Tg cells expressed the activation marker, CD44, to equal extents, (**Figure S5A**) suggesting that MDP administration does not directly influence T cell activation during infection. To address the concern that intraperitoneal administration of MDP may selectively bias T cell accumulation toward the mLN, we repeated the experiment with intravenous administration of MDP, which again resulted in a significant increase in LCMV-specific T cells in the mLN but not the spleen of MDP-treated mice (**Figures S5B and S5C**). To verify that the effect of MDP was NOD2-dependent, we repeated the experiment in WT and littermate *Nod2*^−/−^ mice. NOD2-deficiency resulted in decreased SMARTA-Tg T cell numbers in the mLN (**Figure S5D**) following MDP administration. Furthermore, to mimic the transfer of a more physiological amount of naïve T cells, we observed a similar effect when 10-fold fewer SMARTA cells were transferred^34^ (**Figures S5E and S5F**). Importantly, increased numbers of LCMV-specific T cells in the mLN were independent of transgenic T cell proliferation (**Figures 4J, and S5C**) as well as independent of reduced cell death, since annexin V and propidium iodide staining revealed no differences in the proportions of apoptotic or dead cells between the two groups (**Figure S5G**). Knowing that stromal cells such as fibroblasts provide important survival factors for T cells in SLOs (e.g, IL-7 and IL-15)^35,36^, WT CD4^+^ T cells were co-cultured with mLN stromal cells isolated from WT and *Nod2*^−/−^ mice. We again saw no effect of *Nod2*^−/−^ stroma in maintaining CD4^+^ T cell viability (**Figure S5H**). Since our findings implicate the vascular endothelium in T cell accumulation to the small intestine during infection, we wanted to verify if this effect translated to the mLN. To address this, we infected *Nod2*^*fl/fl*^ or *Nod2*^*ΔCdh5*^ mice with LCMV and MDP administered intraperitoneally, and, three days later, adoptively transferred 5 x 10^5^ congenically-marked (CD45.1) naïve SMARTA-Tg CD4^+^ T cells (**Figure 4K**). Enumerating the amount of transferred SMARTA-Tg cells revealed a BEC-intrinsic role for T cell accumulation in the mLN but not in the spleen (**Figures 4L and 4M**), recapitulating our previous results (**Figures 4F-I and S5D**). Together, these data suggest that NOD2 engagement within BECs leads to increased T cell accumulation in the mLN that is independent of differences in T cell activation, proliferation or cell death.

### Endothelial cell-intrinsic NOD2 engagement increases T cell homing to the mLN

Our data thus far establish a role for BEC-intrinsic NOD2 in mediating T cell accumulation to GALT. We next wanted to address whether NOD2 engagement indeed induces *bona fide* T cell homing to GALT in an endothelial-cell intrinsic way. To answer this question, WT splenocytes labelled with CellTrace™ dye were adoptively transferred into WT and *Nod2*^−/−^ littermates that were pre-treated with MDP 6 days prior and analyzed for the number of labelled splenocytes in recipient mice in the mLN, pLN (auxiliary, brachial and inguinal LNs), and spleen 3 hours post-transfer (**Figure 5A**)^37^. Interestingly, we found that *Nod2*^−/−^ mice had a reduced total number of adoptively transferred T-lymphocytes in their mLNs relative to WT littermates (**Figure 5C**). Interestingly, this effect was not apparent in the spleen, nor in the peripheral lymph nodes (**Figures 5D and 5E**), indicating that NOD2 engagement induces preferential lymphocyte homing to the mLN. To exclude the possibility that lymphocyte-intrinsic NOD2 may contribute to T cell homing to the mLN, we performed a similar assay where fluorescently-labelled, MDP pre-treated WT and *Nod2*^−/−^ splenocytes were adoptively co-transferred at a 1:1 ratio into WT recipient mice (**Figure S6A and S6B**). In these experiments, we found no effect of lymphocyte-intrinsic NOD2 on T cell homing to the mLN and spleen (**Figure S6C and S6D**). We next wanted to see if the effect of NOD2 on T cell homing to the mLN was BEC-intrinsic. To do this, WT splenocytes from congenically-marked CD45.1^+^ C57Bl/6 mice were transferred into recipient CD45.2^+^ *Nod2*^*fl/fl*^ or *Nod2*^*ΔCdh5*^ mice that had been administered MDP 6 days prior, and enumerated the amounts of transferred cells as in **Figures 5A-E**. Our results revealed that BEC-intrinsic NOD2 affects T cell homing specifically to the mLN, as *Nod2*^*ΔCdh5*^ mice had significantly fewer numbers of transferred T cells in their mLNs, but not in their spleen (**Figure 5F and 5G**) relative to *Nod2*^*fl/fl*^ controls. We next examined whether NOD2 affects T cell homing to the mLN during acute LCMV infection by adoptive transfer of splenocytes from SMARTA-Tg mice into WT or *Nod2*^−/−^ littermates infected with LCMV-Armstrong in the presence of MDP 6 days prior (**Figure 5H**). NOD2-dependent homing of SMARTA-Tg lymphocytes was observed into the mLN, but not to the spleen of recipients, 3 hours post transfer (**Figure 5I**). These data cumulatively indicate that BEC-intrinsic NOD2 engagement induces *bona fide* T cell homing to the mLN during homeostasis and infection and provide an explanation for the increased T cells in mLNs of WT mice treated with MDP with or without LCMV (**Figures 4A-I**).

**Figure 5.**
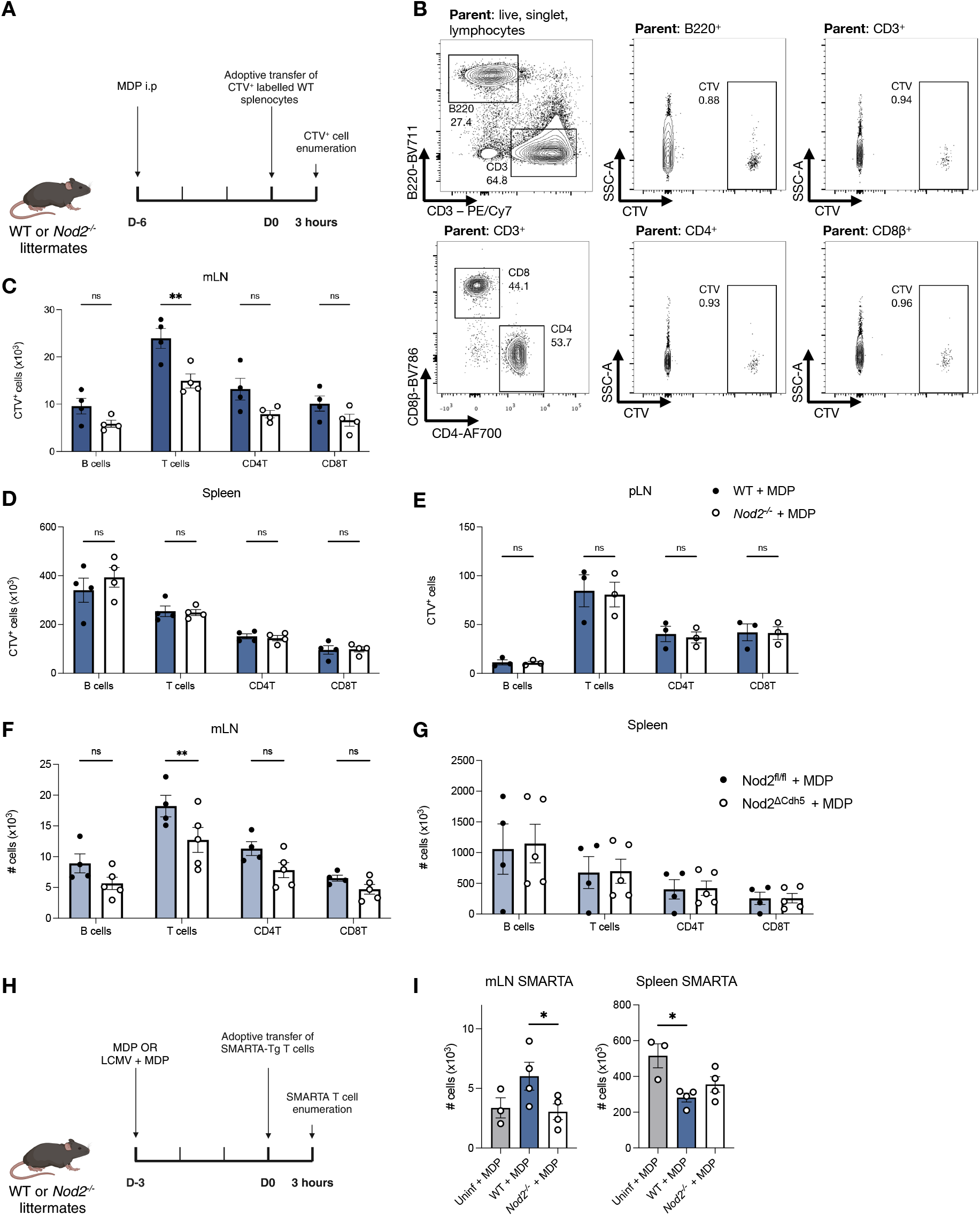
NOD2 engagement leads to *bona fide* T cell homing to the mLN at homeostasis and during infection. (A) Experimental schematic of the *in vivo* homing assay at homeostasis related to panels (C-E). (B) Representative flow cytometry gating strategy of CTV^+^ B cells and T cells. (C-E) Flow cytometric analysis of the number of CellTrace^TM^ violet-positive B and T cell subsets adoptively transferred into recipient WT and *Nod2*^−/−^ mice 3 hours prior. Data show transfer into mLNs (C), spleen (D), and brachial, axial, and auxiliary LNs (E). n = 4 mice per group; ordinary two-way ANOVA. (F-G) Flow cytometry analysis of the number of CD45.1^+^ B and T cell subsets adoptively transferred into recipient CD45.2^+^ *Nod2*^*fl/fl*^ and *Nod2*^*ΔCdh5*^ littermate mice. n = 4-5 mice per group; ordinary two-way ANOVA. (H) Experimental schematic of the *in vivo* homing assay following LCMV-Armstrong infection. Related to panel (I). (I) Flow cytometric analysis of the number of adoptively transferred CD45.1^+^ SMARTA-Tg CD4^+^ T cells into recipient mLNs and spleen. n = 3-4 mice per group; ordinary one-way ANOVA. *p < 0.05; **p < 0.01; ***p < 0.001; ****p < 0.0001. Mean ± SEM depicted.

### NOD2 uniquely synergizes with pro-inflammatory cytokines to induce chemokine and adhesion molecule expression in endothelial cells

We next wanted to establish the ways in which NOD2 intrinsically affects the physiology of endothelial cells following inflammation. Since our previous ABX experiments and scRNA-seq data of mLN BECs, showed an induction of chemokines and adhesion molecules following LCMV infection (**Figures 2H-K and 3D-G**), we focused primarily on the effects of NOD2 engagement on these factors *in vitro*. To do this, BEnd.3 endothelial cells were stimulated with MDP in the presence or absence of TNF and IFNγ, which simulate the Th1-like *ex vivo* inflammation associated with murine LCMV-infection^38^. Expression of chemokines and adhesion molecules, which mediate immune cell recruitment to lymph nodes, were then assessed (**Figures 6A and 6B**). Remarkably, increased expression of *Ccl19* in the presence of TNF and IFNγ was uniquely dependent on NOD2 engagement with MDP (**Figure 6A**), which reflects the NOD2-dependent induction observed following mLN LCMV infection (**Figure 3F**).

**Figure 6.**
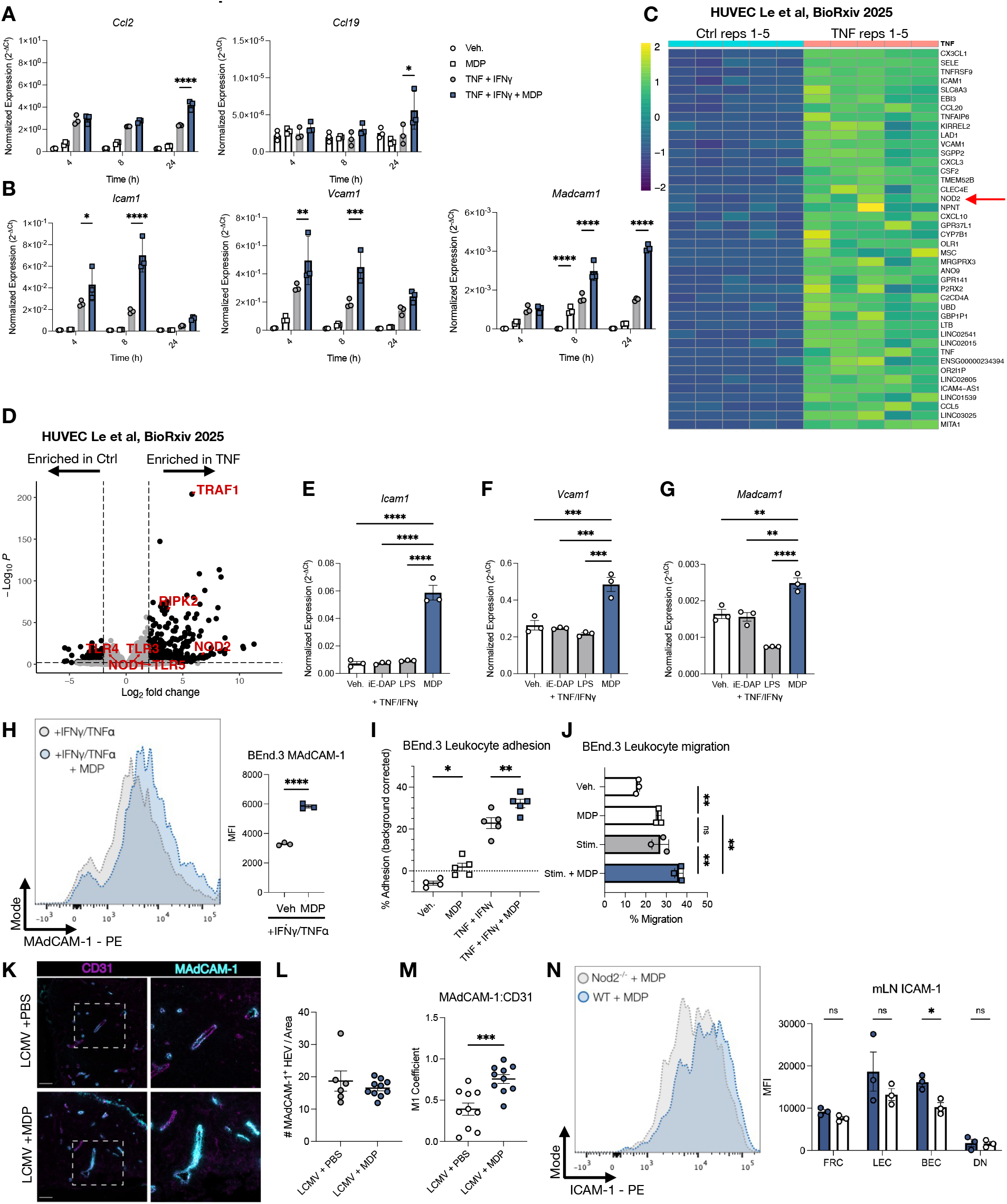
NOD2 uniquely synergizes with pro-inflammatory cytokines to induce chemokine and adhesion molecule expression in endothelial cells. (A-B) Chemokine (A) and adhesion molecule (B) qPCR analysis of BEnd.3 cells treated with or without 20ng/mL IFNγ, 20ng/mL TNF, and 2µg/mL MDP for various times. n = 3 biological replicates per group; ordinary two-way ANOVA. (C) Bulk RNAseq analysis of genes induced by TNF in HUVEC cells. Genes with p_adj_ < 0.0001 and log_2_ fold-change > 6 are plotted. n = 5 replicates. Data are from GSE267418, and generated by Le at al. *BioRxiv*^39^. (D) Bulk RNAseq Volcano plot of indicated PRR and adaptor transcripts induced by TNF in HUVEC cells. N = 5 replicates. Data are from GSE267418, and generated by Le at al. *BioRxiv*^39^. (E-G) qRT-PCR analysis of *Icam1* (E), *Vcam1* (F), and *Madcam1* (G) expression in BEnd.3 cells stimulated with 20ng/mL IFNγ, 20ng/mL TNF, and either 2µg/mL iE-DAP, 2µg/mL MDP, or 0.2µg/mL LPS. n = 3; ordinary one-way ANOVA. (H) Flow cytometry analysis of surface MAdCAM-1 median fluorescence intensity (right) on BEnd.3 cells treated for 24h with IFNγ, TNF, with or without MDP. n = 3 biological replicates per group; unpaired student’s t-test. (I) Splenocyte adhesion assay on BEnd.3 cells treated with indicated treatments for 24 hours. Adhesion percentage is corrected to adhered splenocytes in the absence of BEnd.3 cells. n = 4-5 biological replicates per group; ordinary one-way ANOVA. (J) Splenocyte transmigration assay across BEnd.3 cell monolayers treated with indicated treatments for 24 hours. Stim indicates treatment with 20ng/mL TNF and 20ng/mL IFNγ. n = 3 biological replicates per group; ordinary one-way ANOVA. (K) Confocal microscopy of CD31 and MAdCAM-1 in LCMV-Arm ±MDP infected WT mouse mLNs. (L) Quantification of the total number of CD31^+^ HEVs per image area on 10X objective magnification lens (left images on panel E). Individual data points represent different mouse mLNs. Each point represents one field of view; n = 3 mice per group. (M) Quantification of the Manders’ coefficient of correlation between CD31 and MAdCAM-1 co-expression within mLN HEVs. Individual datapoints represent averages of multiple CD31^+^ HEV correlation calculations within one image. n = 3 mice per group; unpaired student’s t-test. (N) Flow cytometry analysis of surface ICAM-1 expression in WT and *Nod2*^−/−^ mouse mLN stromal cells 6 days following LCMV infection. Median fluorescence intensity is represented across stromal subsets (bottom). n = 3 mice per group; ordinary two-way ANOVA. *p < 0.05; **p < 0.01; ***p < 0.001; ****p < 0.0001. Mean ± SEM depicted.

Furthermore, MDP synergized with TNF and IFNγ to further enhance *Ccl2, Icam1, Vcam1*, and *Madcam1* expression (**Figures 6A and 6B**). Interestingly, MDP alone was sufficient to increase endothelial expression of *Madcam1* to similar levels as TNF and IFNγ priming and further synergized with these cytokines to enhance its expression (**Figure 6B**).

We further wanted to observe if the synergy that exists between TNF/IFNγ and MDP to induce cell adhesion molecule expression was unique among PRRs. We first assessed the effects of inflammation on PRR expression on endothelial cells in a publicly available bulk RNA-seq dataset of TNF-treated human umbilical vein endothelial cells (HUVECs)^39^. Strikingly, following TNF treatment, *NOD2* was among the top 30 most induced genes (**Figure 6C**), and was upregulated to similar extents as canonical BEC cell adhesion molecules (CAMs) such as *ICAM1, VCAM1*, and *SELE*. Furthermore, *NOD2* was uniquely induced among PRRs following TNF treatment (**Figure 6D**), hinting at a unique functional role for NOD2 within the vascular endothelium. To validate these findings, we treated BEnd.3 cells with TNF and IFNγ, in the presence of NOD1, NOD2, and TLR4 ligands (i.e iE-DAP, MDP, and LPS respectively). Strikingly, NOD2 engagement by MDP uniquely synergized with inflammatory cytokines to induce *Icam1, Vcam1*, and *Madcam1* expression (**Figure 6E-G**).

Given the requirement of MAdCAM-1 for T cell homing to GALT, we wanted to confirm that BEC-intrinsic NOD2 engagement leads to heightened endothelial MAdCAM-1 protein expression. MDP administration to BEnd.3 cells induced surface MAdCAM-1 during inflammation (**Figure 6H**). To assess whether NOD2 functionally affects lymphocyte adhesion and migration during inflammation, we observed that BEnd.3 cells stimulated with MDP, TNF and IFNγ increased the adherence (**Figure 6I**) and transmigration (**Figure 6J**) of splenocytes to/across BEnd.3 cells compared to cytokine stimulation alone. We next assessed the ability of NOD2 to regulate mLN adhesion molecule expression *in vivo*. MDP administration in LCMV-Arm-infected mice led to increased MAdCAM-1 expression in mLN BECs (**Figure 6K and 6M**), but did not lead to increases in the relative amounts of HEVs in mouse mLNs (**Figure 6L**). We further observed decreases in ICAM-1 protein expression in mLN BECs of *Nod2*^−/−^ mice relative to WT littermates following infection, (**Figure 6N**), confirming the idea that NOD2 broadly regulates BEC adhesion molecule expression. Taken together, our data implicates NOD2 as a unique PRR within the vascular endothelium that potentiates T cell adhesion and chemoattraction to sites of inflammation.

## Discussion

Aberrant intestinal vascular function is one of the features of Crohn’s disease^40^, and such endothelial dysfunction may result in defects of downstream intestinal immunity. Indeed, the vascular endothelium is required for the spatial coordination of an adaptive immune response following infection^20^; from homing of naïve T cells to secondary lymphoid organs (i.e spleen and lymph nodes) for priming, to directing trafficking to infected tissue for pathogen clearance. Therefore, dysfunctions in the vascular endothelium of gut-associated lymphoid tissue in CD are likely to have downstream effects on effective control of enteric infection and consequent inflammation. Furthermore, CD has a strong genetic component^2^, with mutations in NOD2 being the largest genetic risk factor for CD development^2,3,5^. However, to date, few studies have examined the role of NOD2 in the intestinal endothelium, and how its function within vasculature may affect adaptive immune responses toward infection. Given the idea that aberrancies in intestinal T cell function are associated with CD pathophysiology^41–43^, in our study, we interrogated how NOD2-deficiency affects the vascular endothelium and downstream T cell function in GALT during infection. Our findings demonstrate a requirement for NOD2 in mounting efficient intestinal effector and memory T cell responses following LCMV infection, by modulating the chemokine and CAM expression patterns of blood endothelial cells.

Aberrant T cell population dynamics have been observed in CD and in murine models of IBD. However, much of the literature on T cell dysfunction in CD is confined to cursory examinations of the effector cytokines produced^44^, or on surface protein and transcription factor expression to infer their inflammatory function^42,45^. Such approaches, although informative, oftentimes fail to integrate the overarching concepts of T cell immunity, which involve priming, clonal expansion, and effector and memory specification^46^. Furthermore, the role of CD-associated genetic risk factors in intestinal T cell function is lacking. Naïve T cell homing to the mLNs is a critical initial step for optimal clonal expansion and subsequent T cell responses in the intestine following infection^10,17^. Using LCMV as an enteric pathogen, and as a model for examining T cell expansion, effector, and memory functions^47^, we uncovered a critical role for NOD2 in naïve T cell homing to the mLN during early infection. Furthermore, we find that NOD2 is required for the generation of LCMV-specific effector and memory T cell responses in the ileum, and for protection against secondary viral infection. These data add a further layer of understanding of the role of NOD2 dysfunction in driving CD. For example, although to date CD is thought to be associated with an increase in pro-inflammatory T cell subsets in the intestine that drive inflammation^44^, it remains difficult to decouple whether such increases are induced by inflammation itself, or whether immunological aberrancies existed prior to disease onset. Since our model of LCMV infection allows for the ability to track clonal T cells from the early to late stages of an intestinal immune response, our data suggests that chronic inflammation in CD associated with *NOD2* gene variants may arise from upstream defects in T cell homing to the mLN that occur during early intestinal infections. Furthermore, the inability of NOD2-deficient mice to clear secondary viral infections due to defects in intestinal memory T cell generation suggest that in certain cases of CD, inflammation may be due to defective clearance of recurring intestinal infections. Additionally, our findings are in line with previous work showing that *Nod2*^*- /-*^ animals are more susceptible to the enteric bacterial pathogens *Listeria monocytogenes*^48^, *Citrobacter rodentium*^49^, and *Salmonella typhimurium*^49^, and viral infections such as enteric norovirus^50^, and pulmonary influenza^51^.

Given that NOD2 is a sensor of bacterial MDP, how can NOD2 affect T cell homing to extraintestinal tissue like the mLN in the context of a viral infection? Luminal microbiota-derived MDP has been detected in the circulation of both mice and humans^52,53^ and affects immune responses in extraintestinal tissue^54^. Indeed, microbiota-mediated PAMPs^55^ and metabolites^56^ have been previously shown to regulate immunity in distal sites. We confirm these observations using ABX-depleted mice, which show a requirement for the microbiota in inducing splenic LCMV-specific T cell responses, as well as MAdCAM-1 expression in mLN BECs. Furthermore, we show that mLN MAdCAM-1 expression is rescued upon exogenous administration of MDP. These data suggest that microbiota-derived MDP in the circulation directly affects endothelial functions in the mLN.

A caveat to the antibiotics studies that we performed is that they do not address the question of whether NOD2 engagement within BECs alone is necessary for T cell immunity in the GALT, or whether compensatory mechanisms by other PRRs exist. For example, in other systems, NOD1 or NOD2 stimulation in DCs leads to increases in cross presentation^57^, suggesting the existence of compensatory pathways in NOD signalling. However, our study finds a nonredundant role for BEC-intrinsic NOD2 in mediating vascular endothelial function and downstream T cell immunity in the intestine. Notably, BEC-intrinsic deletion of *Nod2* results in a defect in intestinal memory T cell generation following LCMV infection. This effect likely results from the observed initial defects in T cell homing to mLNs in BEC-intrinsic *Nod2*-deficient mice. Furthermore, we observed that blood endothelial cell induction of CAMs during inflammatory conditions was uniquely dependent on MDP, but not iE-DAP or LPS, suggesting that NOD2 plays a nonredundant role within endothelial cells to mediate adhesion molecule expression and T cell homing during inflammation. These findings offer a novel perspective of the role of NOD2 in CD development. Specifically, low vascular CAM expression within the GALT of CD patients carrying NOD2 gene variants may compromise early immune defense at the level of the intestine, resulting in inefficient clearance of pathogens. This impaired response sets the stage for chronic inflammation as alternative inflammatory pathways become activated. Ultimately, these processes contribute to the chronicity of inflammation associated with CD. Following this logic, insufficient MAdCAM-1 expression in particular may explain why the IBD biologic^58^, Ontamalimab, which is an anti-MAdCAM-1 antibody that blocks the influx of gut-tropic lymphocytes, is ineffective in CD^59^. Future studies providing genetic profiles of individuals with CD failing response to this therapy could determine whether certain polymorphisms make individuals refractory to such treatments. Overall, our work uncovers a previously undescribed role for NOD2 in modulating the vascular endothelium in gut-associated lymphoid tissue, impacting intestinal T cell immunity and control of enteric infection. These studies will stimulate new interest in defining how dysfunctional NOD2 in vascular biology contributes to CD development.

## Acknowledgements

We thank Mary Shi, Christine Pham, and Dr. Troy Ketela at the Princess Margaret Genomics Centre (https://www.pmgenomics.ca/pmgenomics/) for scRNA sequencing processing. We would like to acknowledge Melissa Liu, Dr. Heidi Elsaesser, and Dr. David Brooks for the Vero cells. We would also like to thank Dr. Tania Watts for providing BHK-21 cells, and LCMV-Armstrong and C13 virus stocks. BKT was supported by a Canada Graduate Scholarship, Canadian Association of Gastroenterology Scholarship, and a TRIANGLE Canada Fellowship. This work was also supported by Canadian Institutes of Health Research grants obtained by SEG and DJP.

## Declaration of interests

The authors declare no competing interests.

## Methods

### Animals

All animals were maintained under specific pathogen-free conditions at the University of Toronto’s Department of Comparative Medicine. All experimental animals were littermates, bred from heterozygous crosses. *Nod2*^−/−^ mice were obtained from INSERM, France, *Nod2*^*fl/fl*^ were a gift from Dr. Philip Rosenstiel (Institute of Clinical Molecular Biology, Kiel University), SMARTA-Tg mice were kindly provided by Dr. Tania Watts (University of Toronto). Wildtype and *Cdh5*^*CRE*^ mice were obtained from Jackson Laboratories. All mice were on a C57Bl/6 background, and all experimental animals were maintained on non-acidified water. All experiments were performed between 6-10 weeks of age, and both male and female mice were used. Experiments were conducted as approved by the University of Toronto Animal Care Committee in accordance with the regulations set by the Canadian Council of Animal Care.

### Cell lines

BHK21 cells used for propagation of LCMV were grown in DMEM supplemented with L-glutamine, 10% FBS, Pen/Strep, and 55µM 2-mercaptoethanol. Vero cells used for LCMV PFU measurements were grown in EMEM with 7% FBS and Pen/Strep. BEnd.3 cells (ATCC) used for adhesion assays were grown in DMEM with 10% FBS and Pen/Strep. All cells were grown in an incubator at 37°C and 5% CO2.

### LCMV growth, titration, and storage

LCMV Armstrong and C13 were grown on BHK-21 cells as previously described^60^. Briefly, BHK-21 cells were grown in a T175cm^2^ flask to 70% confluency in high-glucose DMEM + 10% FBS + P/S + 1x 2-mercaptoethanol. Cells were then infected in 10mL of culture media containing virus at an MOI of 0.01 for 1 hour. Subsequently, 35mL of culture medium was added to the flask and cells were incubated with virus for a further 48 hours. Flask supernatant was collected, rid of debris by centrifugation and filtration through a 0.2um filter, and stored at −80°C in 500uL aliquots.

LCMV titration was by infecting Vero E6 cells with pure viral stock or infected tissue homogenate (100mg/mL) for 1 hour at 37°C at various dilutions. Viral supernatant was subsequently aspirated. Plaquing media containing 2X EMEM + 14% FBS + 2% P/S was mixed in a 1:1 ratio with 1% melted ME agarose (Lonza). The media-agarose mixture was overlaid on the infected cells, and the cells were placed in a cell-culture incubator for 6 days. The overlays were then fixed with formaldehyde in PBS and then discarded. The adherent cells were stained with 1% crystal violet. Plaques were enumerated by counting circular regions devoid of crystal violet staining.

### MDP treatments, infections and in vivo SMARTA proliferation assays

For studies assessing the effect of MDP on mLN T cell accumulation and trafficking, at homeostasis, 50µg of MDP (InvivoGen) was intraperitoneally (i.p) injected in mice. Six days later, mouse mLNs were harvested and cell populations were assayed by flow cytometry. For LCMV-infections and assessments of intestinal T cell responses, mice were infected with 2 x 10^5^ PFU of LCMV-Armstrong i.p, and subsequently given 10µg of MDP via i.p injection on days 2, 4, and 6 post-infection. Mice were euthanized at the indicated timepoints in the figure legends. For *in vivo* rechallenge experiments, mice were administered 2 x 10^6^ PFU of LCMV-C13 via intravenous lateral tail vein injection. Twenty-four hours later, mice were euthanized. For *in vivo* SMARTA proliferation experiments, recipient WT C57Bl/6 mice were infected 2 x 10^5^ PFU of LCMV-Armstrong i.p in the presence or absence of 50µg MDP. Splenocytes were isolated from CD45.1^+^ SMARTA-Tg mice, and naïve T cells were isolated using the EasySep™ mouse naïve CD4^+^ T cell isolation kit (Stemcell Technologies) according to manufacturer’s instructions. Naïve CD4^+^ T cells were labelled with 2.5µM CellTrace™ Violet dye (Invitrogen), and 5 x 10^5^ viable labelled T cells were adoptively transferred into recipient mice via lateral tail-vein injection.

### Antibiotics treatment

Mice were administered a cocktail of metronidazole (0.5mg/mL), ampicillin (1mg/mL), neomycin sulphate (1mg/mL), and vancomycin (0.5mg/mL) in 3% sucrose solution in their drinking water for 3 weeks prior to LCMV infection. Sucrose was added to make the antibiotic cocktail more palatable for uptake. Control mice were given the 3% sucrose without the antibiotic cocktail. Mice were infected with LCMV-Armstrong as described and subsequently given PBS or 10µg of MDP via i.p injection every 2 days until the experimental endpoint on day 7. Mice were kept on antibiotics or control sucrose solution for the entirety of the experiment.

### In vivo homing assay

WT splenocytes were harvested in ice cold PBS and RBCs were lysed at room temperature in ACK for 2 minutes prior to being labelled with 1uM CellTrace™ Violet dye (Invitrogen) for 30 minutes. 7.5E6 live labelled cells in 200uL saline were injected intravenously into WT or *Nod2*^−/−^ littermate mice that had been given 50µg MDP intraperitoneally 6 days prior. 3 hours after transfer, recipient mice were euthanized and CellTrace™ positive cells were enumerated in SLOs by flow cytometry. In co-transfer experiments, WT and *Nod2*^−/−^ splenocytes were isolated from littermate mice that had been treated with 50µg MDP 6 days prior. WT and *Nod2*^−/−^ splenocytes were labelled with 1µM CellTrace™ Violet and 1µM CellTrace ™Far Red (Invitrogen) dyes to distinguish between genotypes, prior to resuspending at a 1:1 ratio in PBS. 15E6 cells (7.5E6 cells from each genotype) were injected into WT recipient mice, and homing was allowed to occur for 3 hours prior to SLO harvest.

### T cell survival assays

For *in vivo* T cell survival assays, 5 x 10^5^ naïve SMARTA-Tg CD4^+^ T cells were adoptively transferred into WT mice infected with 2 x 10^5^ PFU of LCMV-Arm with or without 50µg of MDP 3 days prior as in the methods described for SMARTA proliferation assays. Three days following adoptive transfer, isolated mLN lymphocytes were stained with antibodies against CD45.1, CD8β, and CD4 for 20 minutes at 4°C in flow buffer. Cells were subsequently stained with dead cell apoptosis kit with annexin V Alexa Fluor™ 488 & propidium iodide (Invitrogen), and immediately acquired on a BD FACSymphony A3 flow cytometer.

*In vitro* T cell survival assays were performed as previously described^36^. Briefly, mLN stromal cells from WT or *Nod2*^−/−^ mice were isolated (see associated method section). 1E6 stromal cells were plated per well of a 96 well-plate in RPMI 1640 containing 10% FBS, 55µM 2-mercaptoethanol, and Pen/Strep and incubated at 37°C and 5% CO2. After 24 hours, non-adherent cells were removed by washing of the wells three times with PBS, and new media was subsequently added. Adherent stromal cells were allowed to grow for an additional 48h. CD4^+^ T cells were enriched with EasySep™ Mouse CD4^+^ T cell isolation kit according to manufacturer’s instructions (Stemcell Technologies) from SMARTA-Tg spleens. Then, 2 x 10^5^ enriched SMARTA T cells were added on top of the adherent stromal cells and co-cultured for an additional 48 h. Surviving cells were calculated based on the percentage of live cells from the population of CD45.1^+^CD4^+^ cells.

### Lamina propria and intraepithelial lymphocyte isolation

Whole murine small intestines and colons were isolated from mice and cut open longitudinally. Colons were cut into four equal 2cm pieces, whereas the small intestines were cut into 10 2.5cm pieces. Colon pieces were then incubated in 5mL of mucus stripping buffer comprised of Ca^2+^/Mg^2+^-free HBSS (HBSS^-/-^), supplemented with Pen/Strep, 2% FBS, and 5mM dithiothreitol (DTT) for 30 minutes on ice. Small intestines and colons were subsequently incubated in 5mL epithelial stripping buffer (HBSS^-/-^ + 2% FBS, + P/S, + 5mM EDTA) for 15 minutes in a shaking incubator at 37°C. The samples were subsequently vortexed vigorously and the epithelial stripping buffer was subsequently discarded and replaced with a fresh 5mL of buffer. For extraction of IELs, the epithelial fraction was not discarded, but rather passed through a 100µm filter into a 50mL conical tube. This process of epithelial stripping was repeated for a total of three times comprising a total of 45 minutes of shaking incubation at 37°C. IELs were washed with complete RPMI, and stained for flow cytometry analysis. Lamina propria lymphocytes from small intestines and colons were washed for 10 minutes at 37°C with shaking in 10mL of HBSS^-^ 10mM HEPES + P/S, and subsequently digested for 30 minutes at 37°C in 5mL of HBSS^-/-^ + 10mM HEPES + P/S + 0.5mg/mL Collagenase IV + 0.5mg/mL DNAse I. Digestion reactions were stopped with 2mL of complete RPMI (RPMI + 10% FBS + P/S), and samples were passed through a 18x1/2G needle 5 times. Digested samples were purified by passing through 70µm filters over 50mL conical tubes, and samples were then centrifuged at 500xg for 7 minutes. Supernatants were discarded and the resulting cells were purified by means of 40/80% Percoll (Cytiva) gradient centrifugation and used for downstream analyses. For stromal cell analyses, cells were immediately used following digestion and filtration, and Percoll (Cytovia) gradient centrifugation was omitted.

### Lymph node stromal cell isolation

Axial, brachial, inguinal, and mesenteric lymph nodes were isolated and processed separately. Lymph nodes were minced with a scalpel blade on a plastic petri dish and resuspended in 2mL of digestion medium containing RPMI + 2% FBS + 0.8mg/mL Dispase + 0.2mg/mL Collagenase P + 0.1mg/mL DNAse I. Samples were shaken for 15 minutes at 37°C and the samples were vigorously pipetted through a p1000 pipette 20-30x. The turbid supernatants containing stromal cells and leukocytes were collected and passed through 70µm filters into 50mL conical tubes containing PBS supplemented with 5mM EDTA and 2% FBS (flow buffer). The remaining minced lymph node pieces were resuspended in another 2mL of digestion medium and shaken for a further 15 minutes. This process was repeated for a total of three times comprising 45 total minutes of digestion. The resulting cells were centrifuged at 500xg for 5 minutes at 4°C and resuspended in 90uL of flow buffer. 10µL of CD45^+^ Miltenyi microbeads were incubated with the cells for 15 minutes on ice, and stromal cells were isolated using Miltenyi LS columns according to the manufacturer’s instructions. The isolated cells stromal cells were used for downstream flow cytometry analyses.

### Flow cytometry

Leukocytes were extracted from SLOs or lamina propria as described, and dead cells were labelled with Live/Dead™ fixable aqua dead cell stain kit (Invitrogen) in PBS at 4°C for 20 minutes. In experiments using APC gp33-41 class I or PE gp66-77 class II tetramers, tetramer staining was performed in PBS containing 2% FBS, and 0.1% sodium azide (i.e flow buffer) at room temperature for 1 hour prior to viability staining. Lamina propria cells were then surface stained with SuperBright 702 anti-CD19 (1D3) or BV711 anti-B220, APC-efluor780 anti-TCRβ (H57-597), BV786 anti-CD8β (H35-17.2), BUV395 anti-CD4 (GK1.5), SuperBright 600 anti-CD44 (IM7), PerCP-Cy5.5 anti-CD62L (MEL-14), PE-Cy7 anti-CD69 (H1.2F3), BV421 anti-CD103 (M290). Cells were then fixed for 30 minutes at 4°C in IC fixation buffer (eBioscience). For *ex vivo* peptide rechallenge assays, cells were subsequently stained overnight in permeabilization buffer (eBioscience) at 4°C with PE-Cy7 anti-TNFα (MP6-XT22), and BUV737 anti-IFNγ (XMG1.2). FMO controls and tetramer controls were performed where necessary, and stained cells were acquired using a BD FACSymphony flow cytometer (BD Biosciences) and analyzed with FlowJo software (Tree Star, Ashland, OR).

A similar staining protocol was followed for staining mLN and SILP stromal cells, where dead cells were labelled with Live/Dead™ fixable aqua dead cell stain kit (Invitrogen) in PBS at 4°C for 20 minutes. Surface staining was then performed with BUV395 anti-CD45 (30-F11), PE-Cy7 anti-gp38 (8.1.1), BV421 anti-CD31 (390) and primary biotin anti-ICAM-1 (YN1/1.7.4), followed by secondary staining with streptavidin-PE prior to fixation with IC fixation buffer (eBioscience).

### Immunofluorescence microscopy

2-3 mLNs were embedded in optimal cutting temperature (OCT) compound (Sakura Finetek) and placed in 2-methylbutane on dry ice. Embedded samples were frozen at −80°C for a minimum of 24 hours prior to cryosectioning. Cryosections of 7µm were made on a cryostat (Leica Biosystems) Sections were subsequently fixed in −20°C acetone for 15 minutes, and then washed in buffer containing PBS containing 0.2% Tween-20 (PBS-T) three times for 5 minutes at RT. Slides were then blocked in blocking buffer composed of PBS-T + 5% BSA for 1 hour at room temperature. Slides were then incubated in staining buffer composed of PBS-T + 2% FBS with eFluor450 anti-B220 (RA3-6B2), BV421 anti-CD31 (390), APC anti-CD45.1 (A20), purified hamster anti-CD3ε (500A2), and biotin anti-MAdCAM-1 (MECA-367) at overnight at 4°C. Slides were washed 3x in PBS-T for 5 minutes, prior to staining with AF594 anti-hamster, and PE streptavidin antibodies for 1 hour at room temperature. Slides were again washed 3x in PBS-T for 5 minutes, and, where applicable, stained with Hoechst 33342 nucleic acid stain (Invitrogen) for 20 minutes. Slides were mounted with Dako fluorescent mounting medium (Agilent). Images were taken on a Zeiss LSM700 confocal microscope using the Zen software. Cell counts were quantified using the CellProfiler^61^ software, whereas Manders’ coefficients were calculated using the JACoP^62^ plugin on ImageJ.

### Ex vivo lamina propria peptide rechallenge

Isolated SILP leukocytes from mice infected with LCMV-Armstrong 30 days prior were stimulated for 4h with 5µg/mL of the MHC Class II-restricted LCMV gp61-80 (Anaspec), or 2µg/mL of the MHC Class I-restricted LCMV gp33-41 (Genscript), in the presence of 20ng/mL IL-2 (Peprotech) and protein transport inhibitor cocktail (eBioscience). Cells were then stained for viability, surface markers, and intracellular cytokines as indicated in the flow cytometry section.

### In vivo LCMV-C13 rechallenge and cytokine stain

Mice infected with 2 x 10^5^ PFU LCMV-Armstrong i.p 30 days prior were infected with 2 x 10^6^ PFU LCMV-C13 intravenously. Twenty-four hours later, SILP leukocytes were isolated and first stained with Live/Dead™ fixable aqua dead cell stain kit (Invitrogen), and then with fluorescent antibodies against TCRβ, CD4, CD8β, B220, CD44, IFNγ and TNF. Staining with IFNγ and TNF was done overnight in permeabilization buffer at 4°C, prior to running on a flow cytometer. Importantly, no *ex vivo* treatment with protein transport inhibitor cocktails was performed, in order to capture cells that are uniquely responding to LCMV-C13 as done previously^63^.

### BEnd.3 cell treatments and adhesion assays

For gene expression analysis confluent BEnd.3 cells (ATCC) from a T75cm^2^ flask were seeded into 24 wells of 6 well plates. BEnd.3 cells were grown for 24 hours until they reached 80% confluency, and were subsequently treated with 2µg MDP, or 2µg iE-DAP, or 0.2µg LPS with or without 20ng/mL TNFα and 20ng/mL IFN*γ* in 1mL of used cell media (DMEM + 10% FBS + P/S) for various time periods. Cells were subsequently lysed in RLT buffer for RT-qPCR gene expression analysis.

For adhesion assays, 5E4 BEnd.3 cells were seeded into 24 well plates, and grown for 48 hours until 80-90% confluence. Cells were treated with 2µg MDP with or without 20ng TNFα and 20ng IFN*γ* in 500uL of used cell media for 18 hours. Media was aspirated and 5 x 10^5^ WT splenocytes resuspended in basal RPMI were added to each well of BEnd.3 cells and allowed to adhere for 30 minutes at 37°C with shaking at 80rpm. Nonadherent cells were removed by two washes with PBS, and the remaining adherent cells were lifted mechanically by vigorous pipetting and subsequently analyzed by flow cytometry.

### Transendothelial leukocyte migration assay

BEnd.3 cells were grown in T75cm^2^ flask to 80% confluency in complete DMEM. BEnd.3 cells were trypsinized and resuspended to a concentration of 4 x 10^5^ cells/mL in complete DMEM.

100µL of cell mixture was pipetted on top of 5µm transwells (Corning) in 24 well plates. 600µL of complete DMEM was added to the bottom chamber, and cells were allowed to grow for 24 hours. BEnd.3 cells were then treated with 2µg/mL MDP, and/or 20ng/mL TNFα and 20ng/mL IFN*γ* in 100µL complete DMEM for another 24 hours. Before the migration assay, excess media in the bottom chamber was removed, and replaced with fresh migration medium (basal RPMI-1640, Wisent) supplemented with 200ng/mL murine CCL21 (Peprotech). Excess treatment medium on the transwells was removed. WT splenocytes were isolated and diluted to 6E6 cells/mL in migration medium, and 100µL of cell suspension was added on top of the BEnd.3 cells. Cells were allowed to migrate for 4 hours at 37°C. Migrated cells in the bottom chamber were harvested and counted on a hemocytometer.

### RNA extractions and real-time quantitative polymerase chain reactions

For tissue-based RNA extractions, entire lymph nodes, 100mg spleen, 2cm of proximal colon, or 2cm of terminal ileum were placed in 650µL buffer RLT supplemented with 2-mercaptoethanol, and RNA was extracted according to RNeasy Mini kit instructions (Qiagen). RNA quality and quantity was measured on a Nanodrop spectrophotometer (ThermoFisher Scientific). Samples with A260/280 ratios lower than 1.9, and/or A260/230 ratios lower than 1.5 were not used. RNA concentration was normalized to 200ng/µL and reverse transcription was performed on 2µg of RNA (10µL) using High-capacity cDNA reverse transcription kit (Applied Biosystems) according to manufacturer’s instructions. cDNA was diluted 1/10 and qRT-PCR was performed using PowerUp™ SYBR™ Green master mix (ThermoFisher Scientific) and raw Ct values were obtained from a CFX384 TouchTM Real-Time PCR Detection System (Bio-Rad). Melt curve analysis was performed to verify primer specificity, and gene expression was normalized to that of the housekeeping gene Rpl19. The same process was performed for qPCR analysis of gene expression data from BEnd.3 cells (ATCC) seeded in 6 well plates, apart from lysis in 350µL of buffer RLT in this case. Primer sequences are listed in Table S1.

### Single-cell RNA sequencing

Eight WT mice and 8 *Nod2*^−/−^ mice were randomized into two groups: uninfected + MDP, and infected + MDP, such that 4 WT and 4 *Nod2*^−/−^ mice received either 50µg of MDP i.p alone, or 50µg of MDP + 2 x 10^5^ PFU LCMV-Armstrong i.p. Six days post treatment, the entire mLN chain stroma was isolated using magnetic enrichment as described. mLN stromal cells from each individual treatment were pooled together, and cells were fixed and permeabilized, and stored at −80°C in a glycerol solution for 1 week prior to 10X Chromium Fixed RNA Profiling (10X genomics) according to the long-term storage fixation protocol provided by 10X Genomics.

Quality assurance for sample viability was performed via trypan blue and automatic cell counting to ensure that all samples had a minimum viability of 70%. 10X Chromium mouse transcriptome probes including an eGFP spike (Chromium Mouse Transcriptome Probe Set) were hybridized to the sample mRNA, and additionally filtered through a 20µm filter to remove cell clumps according to the manufacturer’s protocol. The hybridized samples were then partitioned into barcoded gel beads-in-emulsion (GEMs) on a microfluidic chip prior to sequencing on a Novaseq X plus platform (Illumina) with 28/10/10/90 base pair cycles at the Princess Margaret Genomics Centre at the University Health Network. CellRanger (version 7.2.0, 10X Genomics) was used to align reads to the mm10-eGFP-2020-A transcriptome and perform quality control. Filtered feature matrices were used for downstream analysis in R.

### Single-cell RNA sequencing analysis

The single-cell RNA sequencing data from baseline and infected WT and *Nod2*^−/−^ mLNs was analysed with R (version 4.3.3) and Seurat (version 5)^64^. Quality control excluded genes that were not present in at least 6 cells, or cells with < 600 genes detected, or 10% mitochondrial gene expression. Immune cells were removed by removing cells with *Ptprc* expression. Datasets were merged, normalized, and scaled using the SCTransform function. An elbow plot was used for the selection of principal components. After dimensionality reduction, the FindNeighbors and FindClusters (resolution = 0.4) functions were used for determining the amount of cell subsets.

To identify the most highly differentially expressed genes across clusters, Seurat’s FindConservedMarkers function was used. Clusters were manually annotated based on previously identified stromal marker expression^65–68^. Differential expression within clusters was performed using the FindMarkers function. Volcano plots and gene set enrichment analyses were performed with the EnhancedVolcano and clusterProfiler^69^ packages respectively. All associated sequencing files and processed Seurat object will be available on the gene expression omnibus as record GSE298513.

### Analysis of publically available scRNA-seq and bulk RNA-seq data

For scRNA-seq, pediatric, fetal, and adult endothelial cell normalized data was obtained from the Space-Time Gut Cell Atlas at gutcellatlas.com^28^. Endothelial cells were visualized using the uniform manifold approximation and projection (UMAP) method. For bulk RNA-seq data of HUVEC cells^39^, data was accessed through Gene Expression Omnibus (GEO) GSE267418. Differential gene expression between CTRL1_EGM_REP1-5 and HIT1_TNF_REP-15 was performed using DESeq2.

#### Quantification and statistical analyses

Prism software (Graphpad v10.4.1) was used for all statistical analyses. All statistical tests performed were either unpaired two-tailed t-tests, one-way ANOVA, or two-way ANOVA depending on the amount of variables present in the dataset. Statistical tests for multiple comparisons were performed where applicable. Statistical significance was defined as *p ≤ 0.05, **p ≤ 0.01, ***p ≤ 0.001, ****p ≤ 0.0001. Results are presented as mean ± standard error mean (SEM).

---

**Supplementary table S1.**
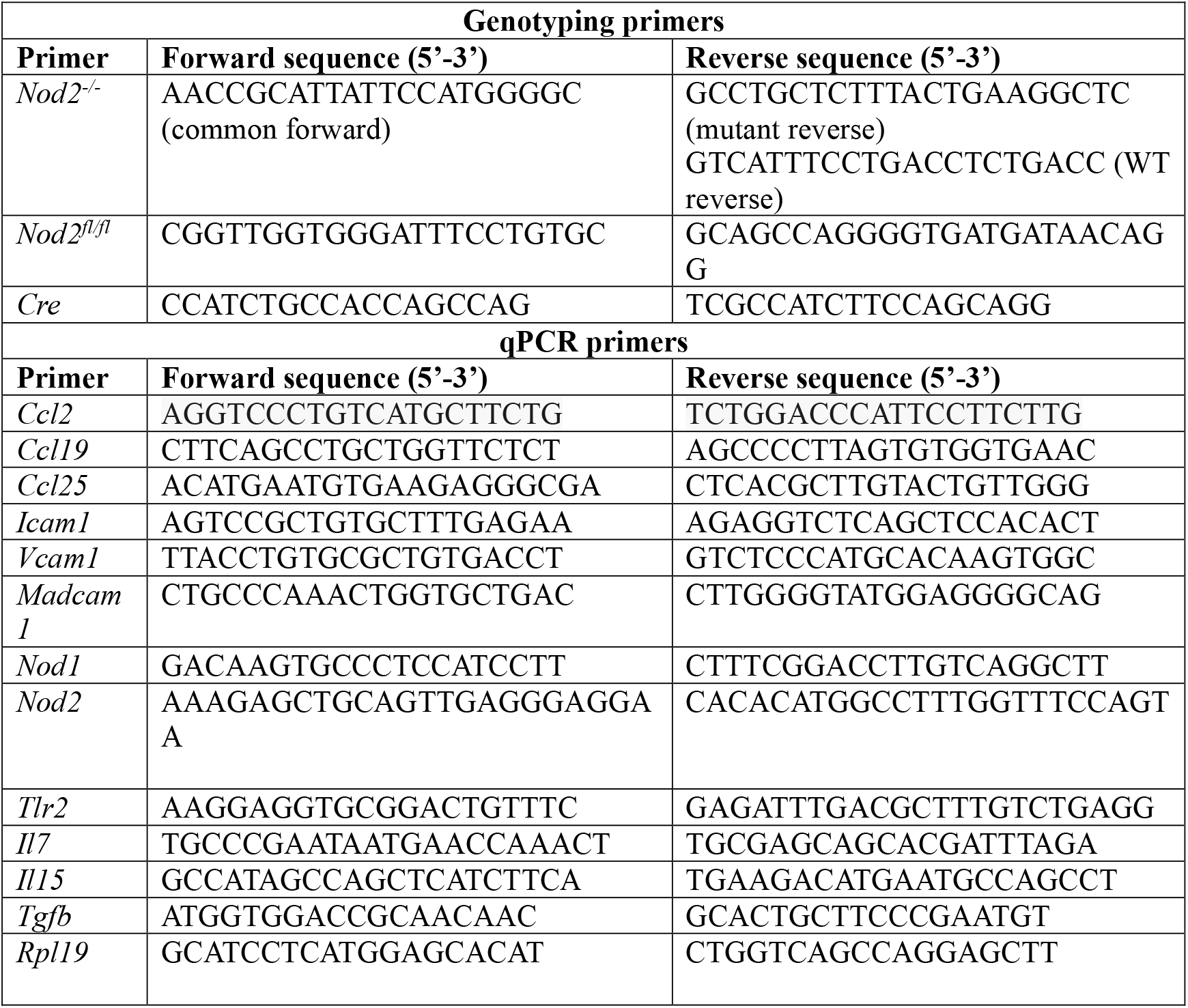
Primers used in this study.

## Notes

### Competing Interest Statement

The authors have declared no competing interest.

