## Supplemental Figures for "Endothelial cell-intrinsic NOD2 signaling regulates the intestinal immune response through the generation of effector and memory T cells"

**Figure S1 – NOD2-mediated enhancement of effector and memory T cell responses are SILP specific, and not due to increases in intestinal chemokines or T cell maintenance cytokines. Related to Figure 1.**

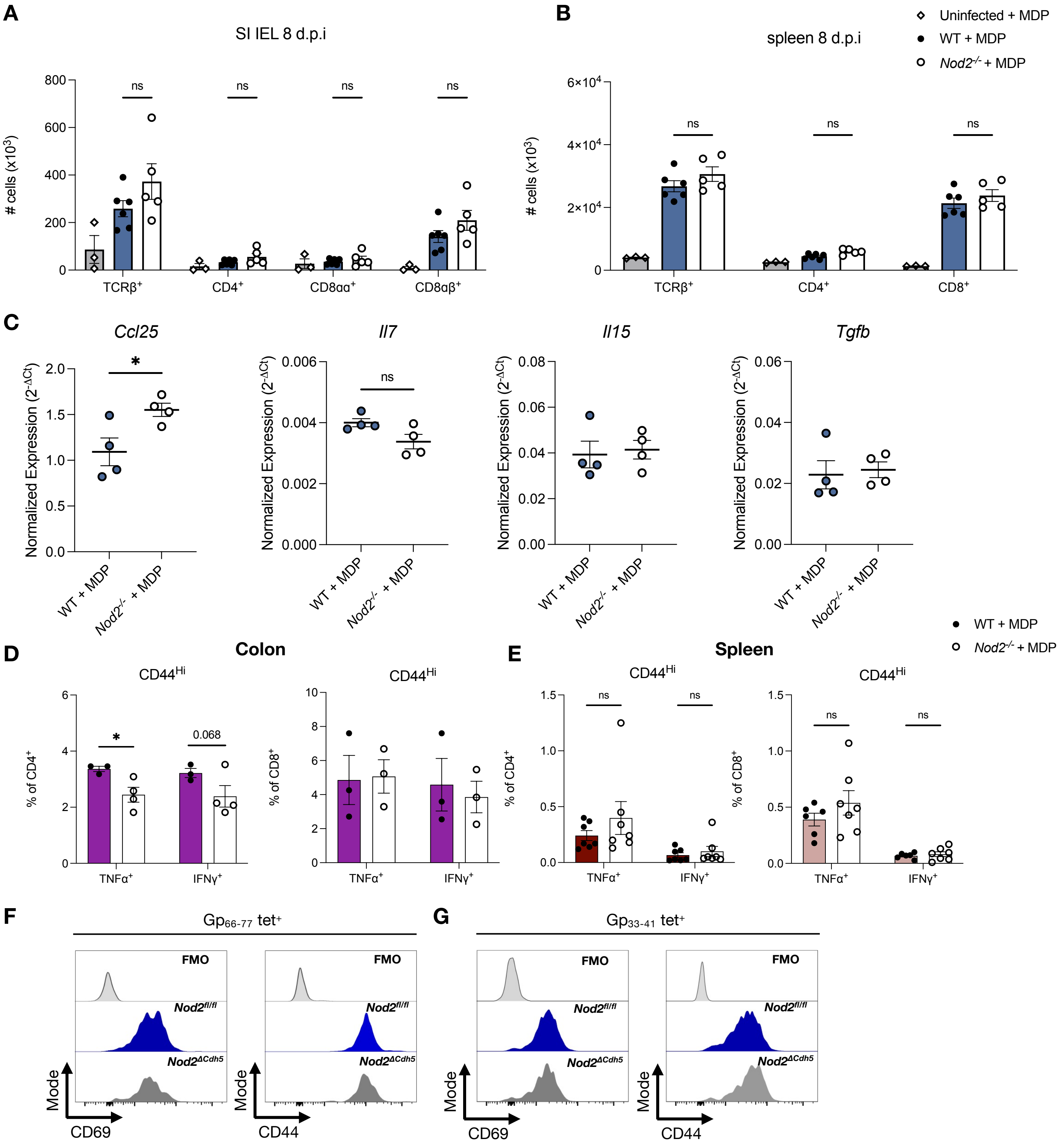

**Figure S1. NOD2-mediated enhancement of effector and memory T cell responses are SILP specific, and not due to increases in intestinal chemokines or T cell maintenance cytokines. Related to Figure 1.**
(A) Small intestinal intraepithelial lymphocyte number enumeration by flow cytometry 8 days post LCMV-Armstrong infection. n >= 3 mice per group; ordinary two-way ANOVA.
(B) Splenic lymphocyte number enumeration by flow cytometry 8 days post LCMV-Armstrong infection. n >= 3 mice per group; ordinary two-way ANOVA.
(C) qRT-PCR analysis of *Ccl25, Il7, Il15,* and *Tgfb* mRNA transcripts in the terminal ileum 8 days post infection. n = 4 mice per group; unpaired student's t-test.
(D, E) *Ex vivo* cognate peptide restimulation assays of colonic (D), and splenic (E) $CD4^+$ (top) and $CD8^+$ (bottom) 30 days following LCMV-Armstrong infection. n >= 3 mice per group; ordinary Two-way ANOVA.
(F-G) Representative histograms of $CD44^+$ and $CD69^+$ marker expression of $gp_{66-77}$ tetramer$^+$ $CD4^+$ (F) and $gp_{33-41}$ tetramer$^+$ $CD8^+$ (G) T cells from *Nod2*$^{fl/fl}$ and *Nod2*$^{\Delta Cdh5}$ mice.
*p < 0.05; **p < 0.01; ***p < 0.001; ****p < 0.0001. Mean ± SEM depicted.

### Figure S2 – High NOD2 expression within mLN BECs. Related to Figure 3.

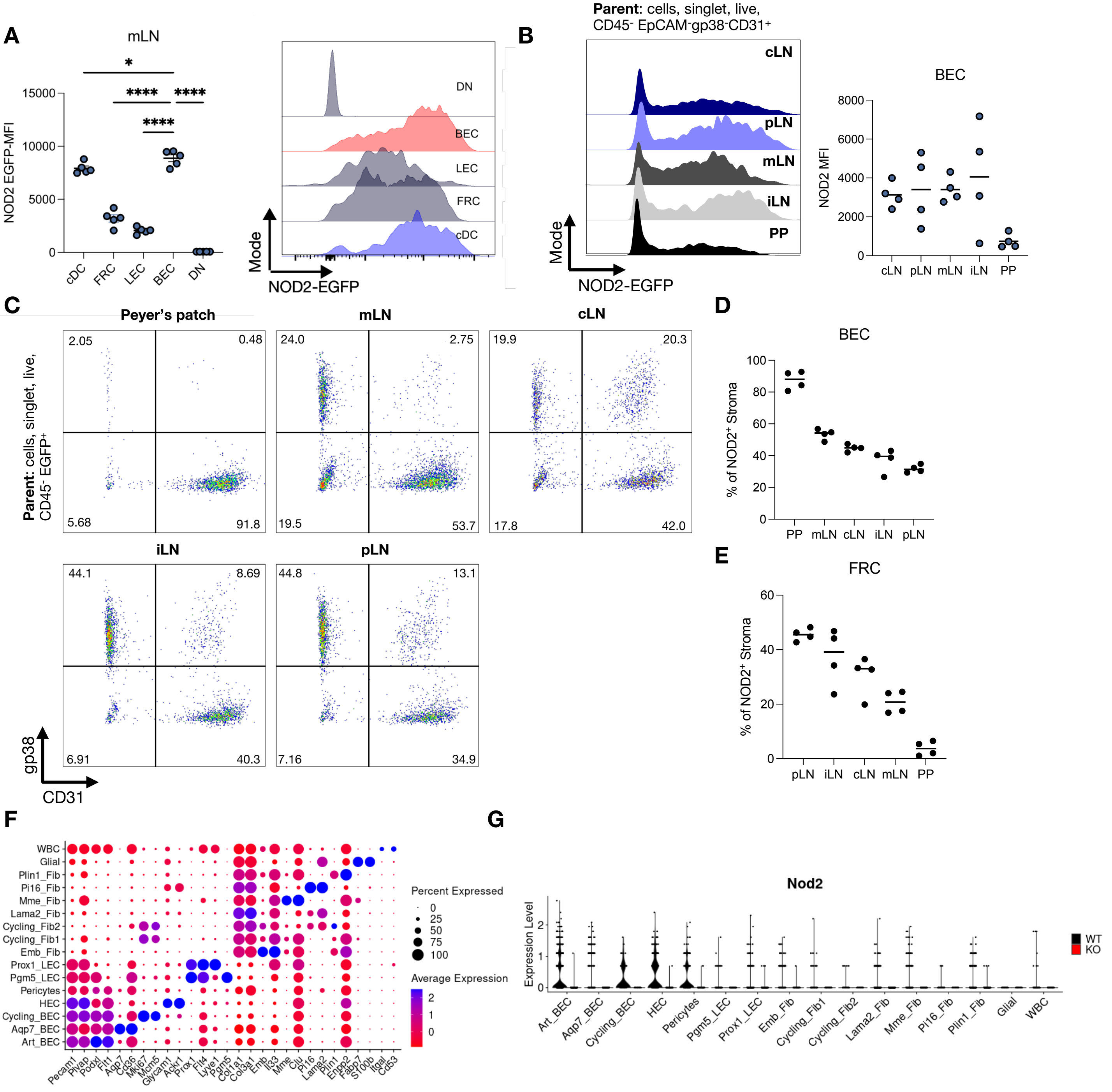

**Figure S2. High NOD2 expression within mLN BECs. Related to Figure 3.**
(A) Flow cytometric analysis of NOD2-EGFP expression in cDCs and stromal cell subsets within naïve mLNs. n = 5 mice; ordinary one-way ANOVA.
(B) Flow cytometric analysis of NOD2-EGFP expression within cervical (cLN), peripheral (pLN), mesenteric (mLN), iguinal (iLN) lymph nodes and Peyer's patches (PP) of naïve mice. n = 4 mice per group.
(C-E) Flow cytometric analysis of the proportion of BECs (D) or FRCs (E) among NOD2-EGFP expressing stromal cells ($CD45^-$). n = 4 mice per group.
(F) Expression dot plot of marker genes used to classify all cell clusters within NT and LCMV datasets. Blood endothelial cell (BEC), high endothelial cell (HEC), lymphatic endothelial cell (LEC), fibroblast (Fib), white blood cell (WBC).
(G) Violin plot quantification of *Nod2* expression within the indicated cell types. Data are aggregated from both the NT and LCMV WT and $Nod2^{-/-}$ mouse datasets.
$^*p < 0.05$; $^{**}p < 0.01$; $^{***}p < 0.001$; $^{****}p < 0.0001$. Mean ± SEM depicted.

**Figure S3 – NOD2 engagement does not affect mLN stromal populations and gene expression of fibroblasts and LECs during LCMV infection . Related to Figure 3.**

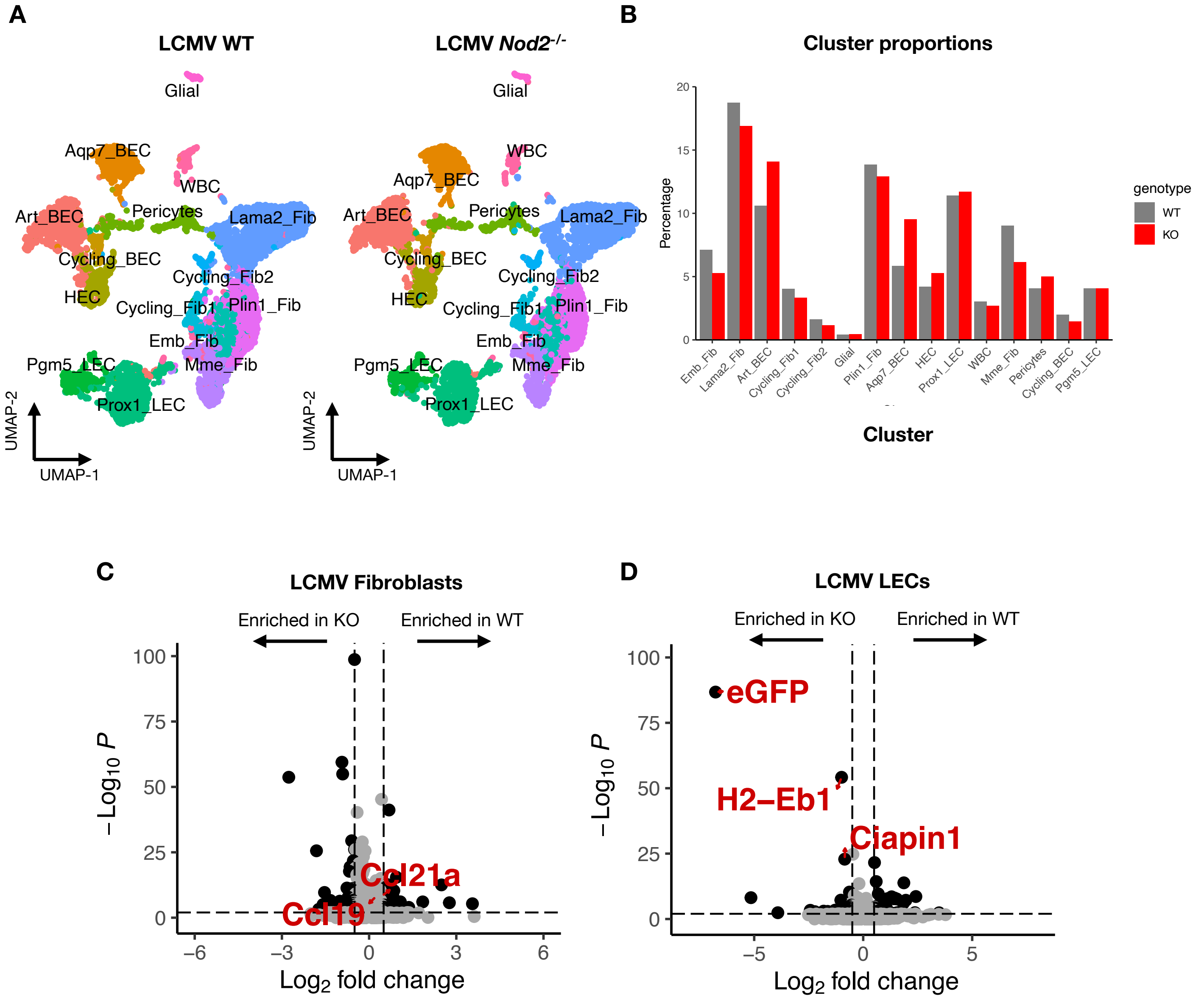

**Figure S3. NOD2 engagement does not affect mLN stromal populations and gene expression of fibroblasts and LECs during LCMV infection. Related to Figure 3.**

(A) scRNA-seq UMAP plot of mLN stromal cells from LCMV-infected (6 days) WT and *Nod2*$^{-/-}$ littermates.

(B) Quantification of the cell population number of each UMAP cluster of WT and *Nod2*$^{-/-}$ LCMV-infected mLNs.

(C-D) Volcano plots of the DGE analysis between LCMV-infected WT and *Nod2*$^{-/-}$ mLN FRCs (C) and LECs (D).

**Figure S4 – NOD2 deficiency does not affect iLN, splenic, and small intestinal lamina propria T cell populations at baseline. Related to Figure 4.**

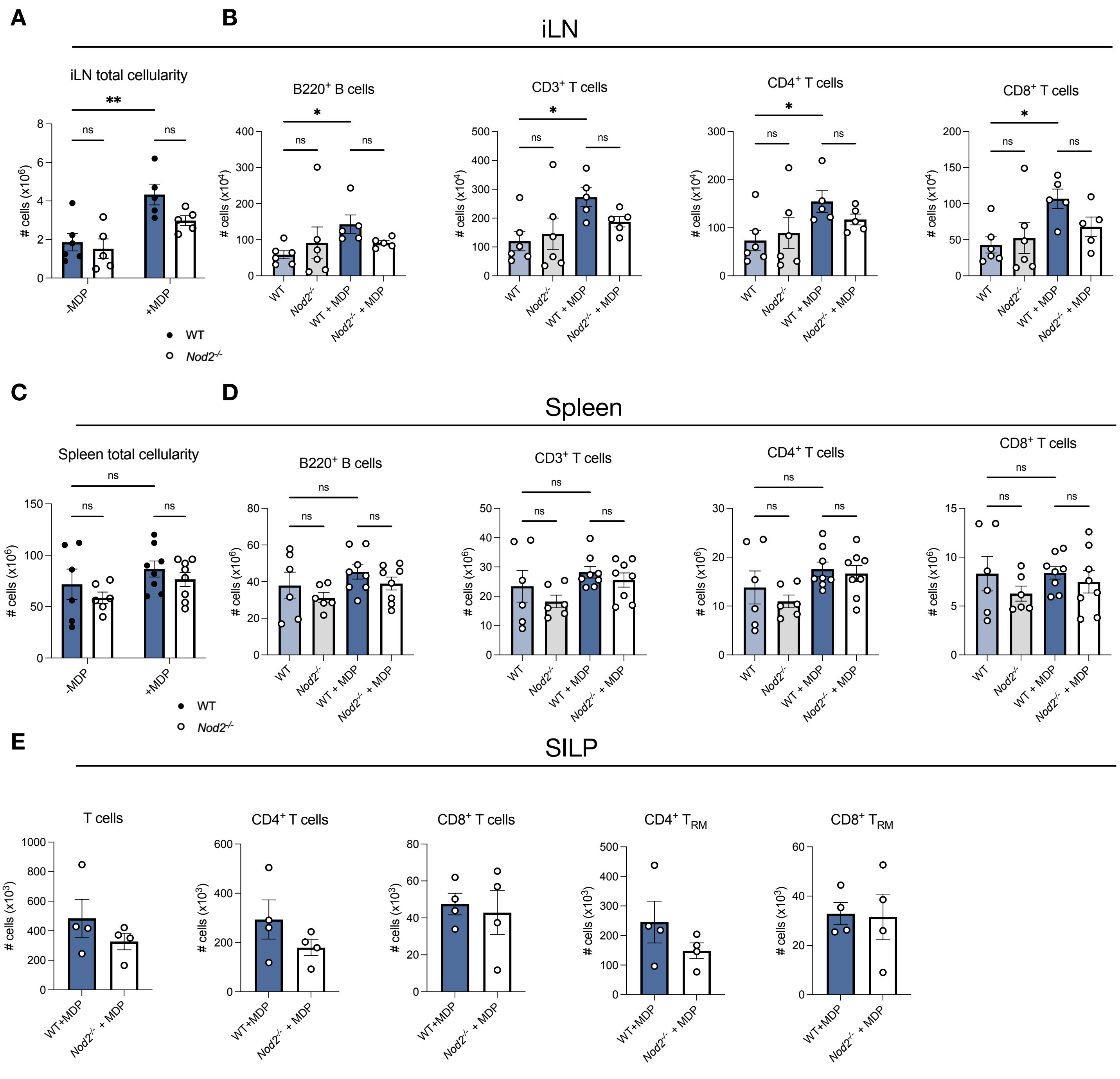

**Figure S4. NOD2 deficiency does not affect iLN, splenic, and small intestinal lamina propria T cell populations at baseline. Related to Figure 4.**
(A, C) Total live leukocyte cellularity of the iLN (A) and spleen (C) of WT and *Nod2*$^{-/-}$ littermates. n = 4-5 mice per group; ordinary two-way ANOVA.
(B, D) Flow cytometry analysis of the number of B and T cell subsets in the iLN (B) and spleen (D) of WT and *Nod2*$^{-/-}$ littermates. n = 4-5 mice per group; ordinary one-way ANOVA.
(E) Flow cytometry analysis of bulk $CD4^+$, $CD8^+$, $CD4^+CD62L^-CD44^+CD69^+$ ($CD4^+$ $T_{RM}$), or $CD8^+CD62L^-CD44^+CD69^+$ ($CD8^+$ $T_{RM}$) in the SILP of WT and *Nod2*$^{-/-}$ littermates. n = 4 mice per group; unpaired student's t-test.
*p < 0.05; **p < 0.01; ***p < 0.001; ****p < 0.0001. Mean ± SEM depicted.

**Figure S5 – Enhanced T cell accumulation in the mLN upon MDP administration is independent of the route of administration and the amount of adoptively transferred cells. Related to Figure 4.**

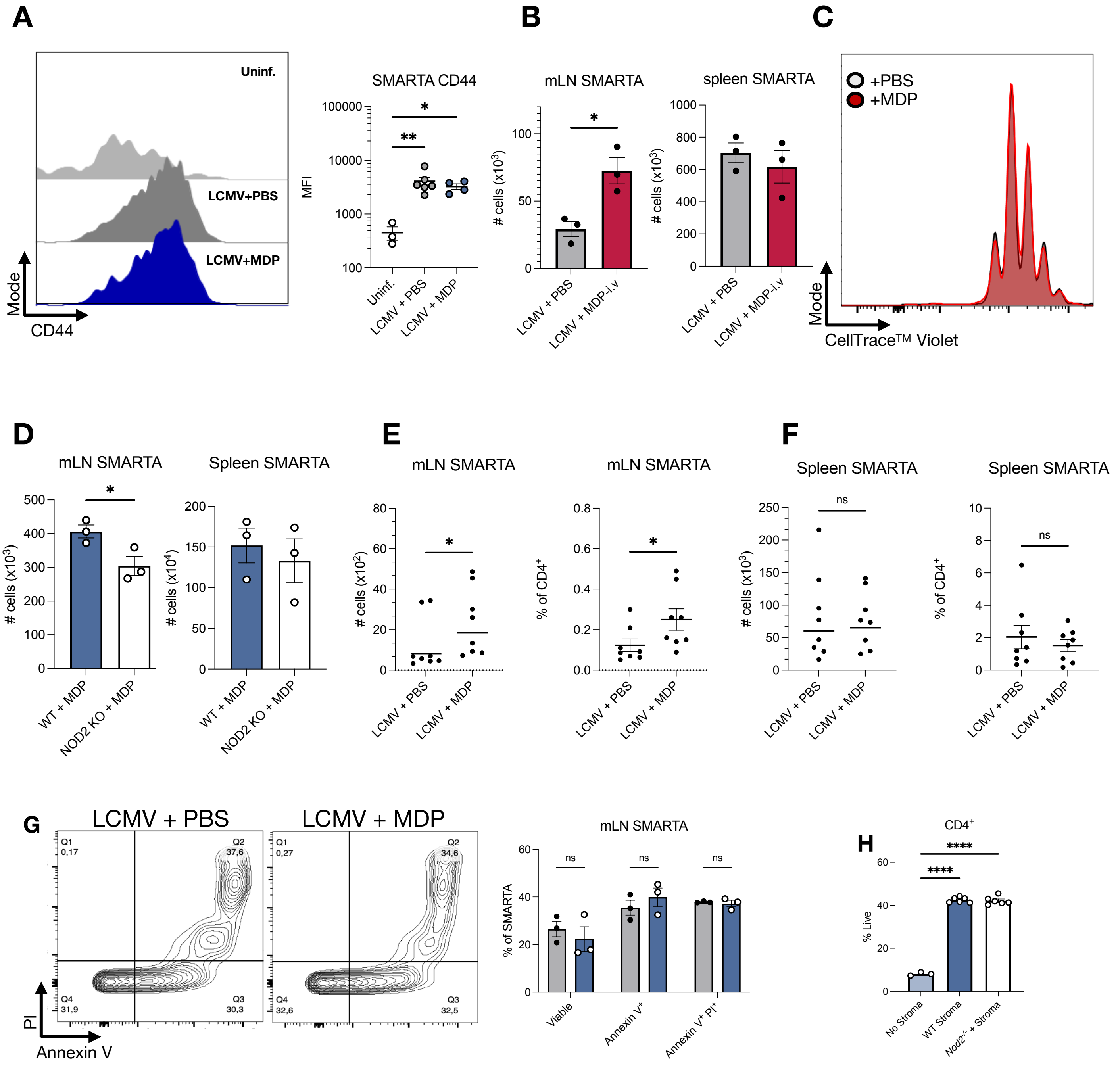

**Figure S5. Enhanced T cell accumulation in the mLN upon MDP administration is independent of the route of administration and the amount of adoptively transferred cells. Related to Figure 4.**

(A) Characterization of surface CD44 expression on SMARTA-Tg T cells 72-hours post adoptive transfer. n > 3 mice per group; ordinary one-way ANOVA.

(B) Flow cytometry analysis of the number of adoptively transferred SMARTA-Tg T cells in the mLN (left) and spleen (right) of LCMV-infected mice with or without intravenous (i.v) MDP administration. Mice were euthanized 72 hours post adoptive transfer. n = 3 mice per group; unpaired student's t-test.

(C) CellTrace™ violet proliferation assay of the adoptively transferred SMARTA-Tg T cells in the mLN of infected recipient mice. n = 3 mice per group.

(D) Flow cytometry analysis of the number of adoptively transferred SMARTA-Tg T cells in the mLN and spleen of LCMV-infected WT and *Nod2*$^{-/-}$ littermate mice. n = 3 mice per group; unpaired student's t-test.

(E, F) Flow cytometry analysis of the number of adoptively transferred SMARTA-Tg T cells in the mLN (E) and spleen (F), 3 days following transfer. 50,000 SMARTA-Tg cells were transferred in this experiment, whereas 500,000 cells were transferred in panels (A-C). n = 8 mice per group; unpaired student's t-test.

(G) Annexin V & propidium iodide viability stain of adoptively transferred CD45.1 SMARTA-Tg $CD4^+$ T cells 3 days post-adoptive transfer. n = 3 mice per group; ordinary two-way ANOVA.

(H) *Ex vivo* SMARTA-Tg $CD4^+$ T cell viability 48 hours post plating on stromal cells isolated from WT or *Nod2*$^{-/-}$ mLNs.

$*p < 0.05$; $**p < 0.01$; $***p < 0.001$; $****p < 0.0001$. Mean ± SEM depicted.

**Figure S6 – T cell-intrinsic NOD2 is dispensable for mLN homing. Related to Figure 5.**

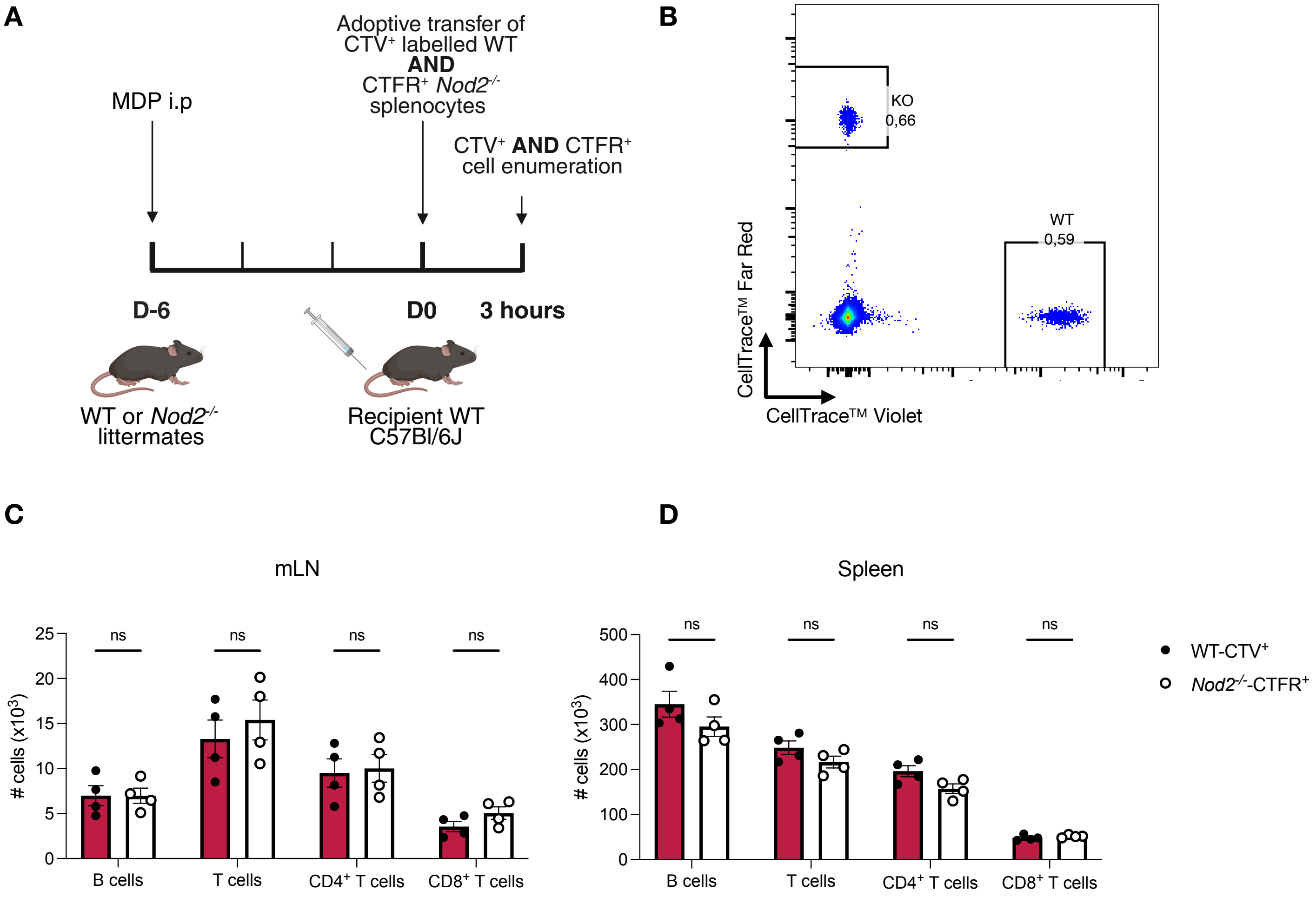

**Figure S6. T cell-intrinsic NOD2 is dispensable for mLN homing. Related to Figure 5.**
(A) Experimental schematic of competitive *in vivo* homing assays.
(B) Representative flow plot showing 1:1 input of WT and *Nod2*$^{-/-}$ T cells.
(C-D) Enumeration of transferred lymphocytes into the mLN (C) and spleen (D) of WT and *Nod2*$^{-/-}$ recipients by flow cytometry. n = 4 mice per group; ordinary two-way ANOVA.
Mean ± SEM depicted.

#### Gating strategy for lamina propria T cells

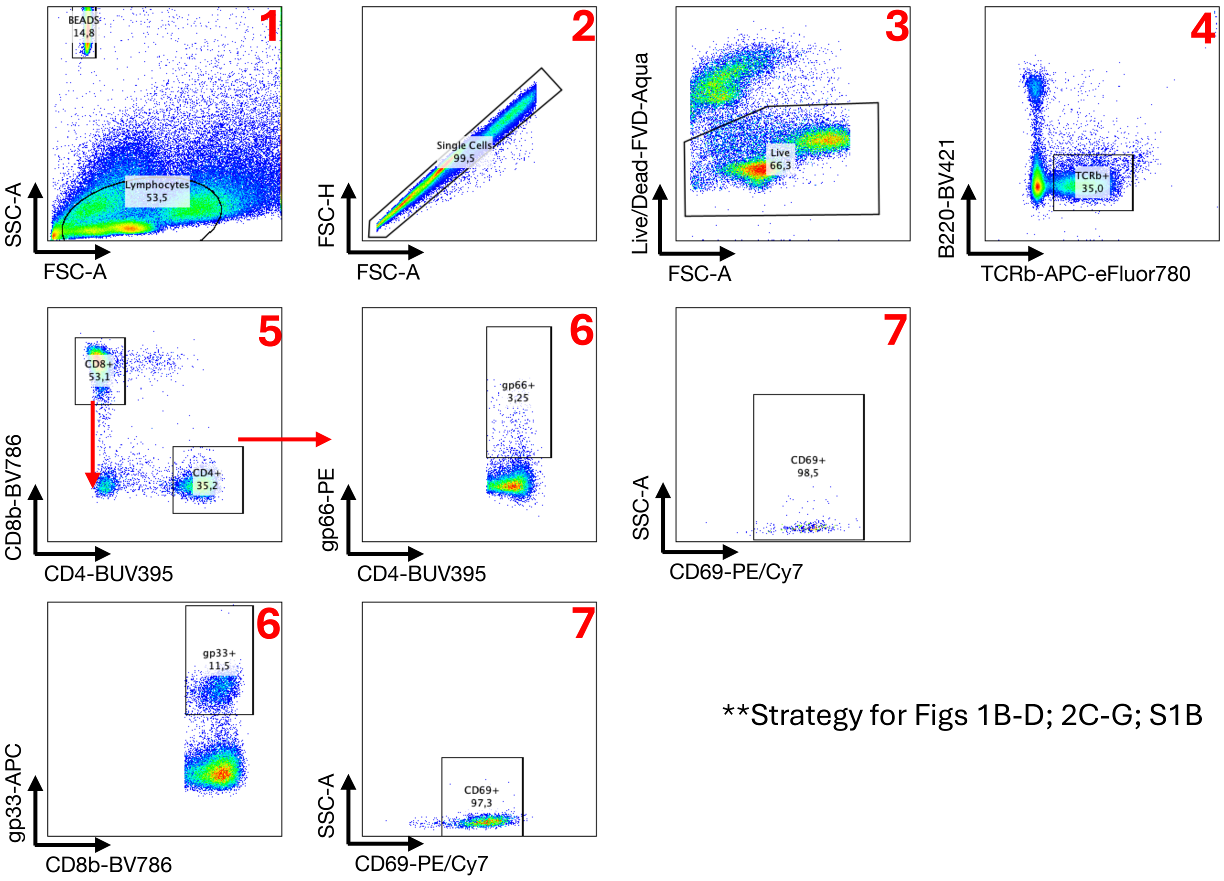


**Strategy for Figs 1B-D; 2C-G; S1B

#### Gating strategy for IEL T cells

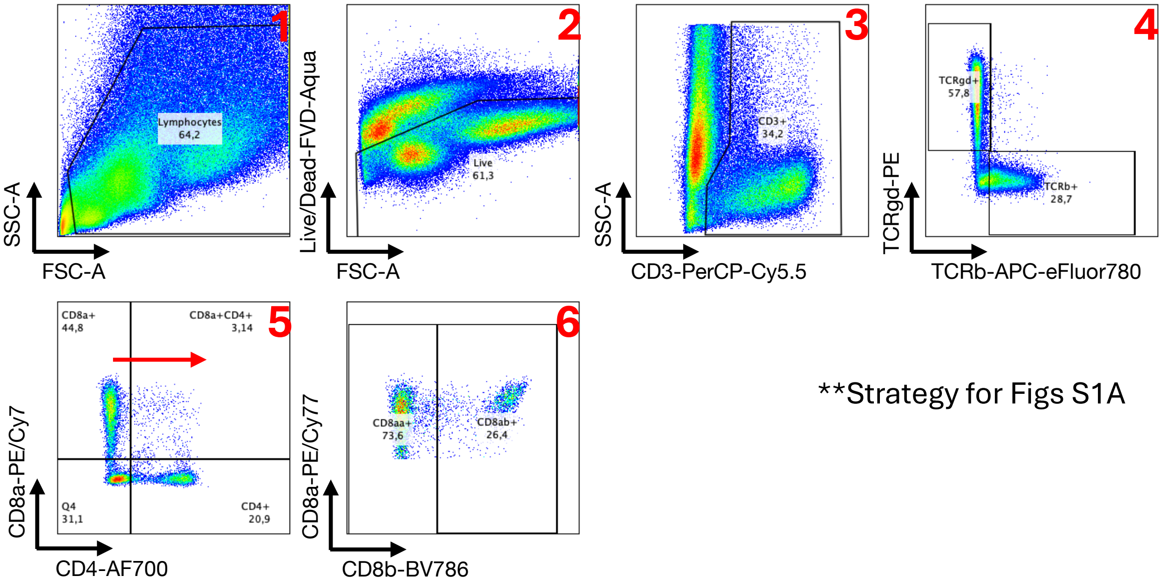


**Strategy for Figs S1A

#### Gating strategy for lamina propria T cell *ex vivo* peptide restim

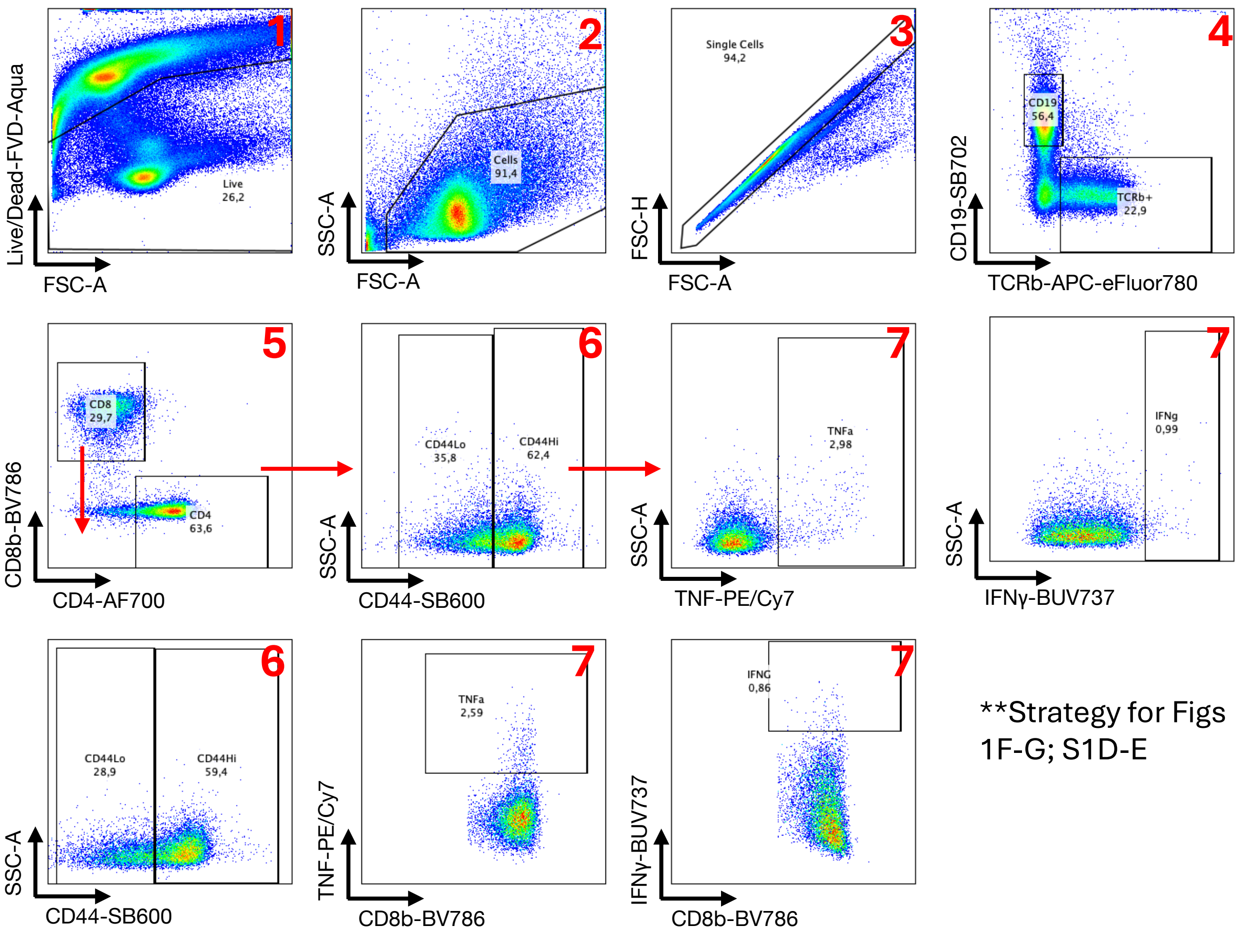


#### Gating strategy for lamina propria *in vivo* LCMV C13 restim

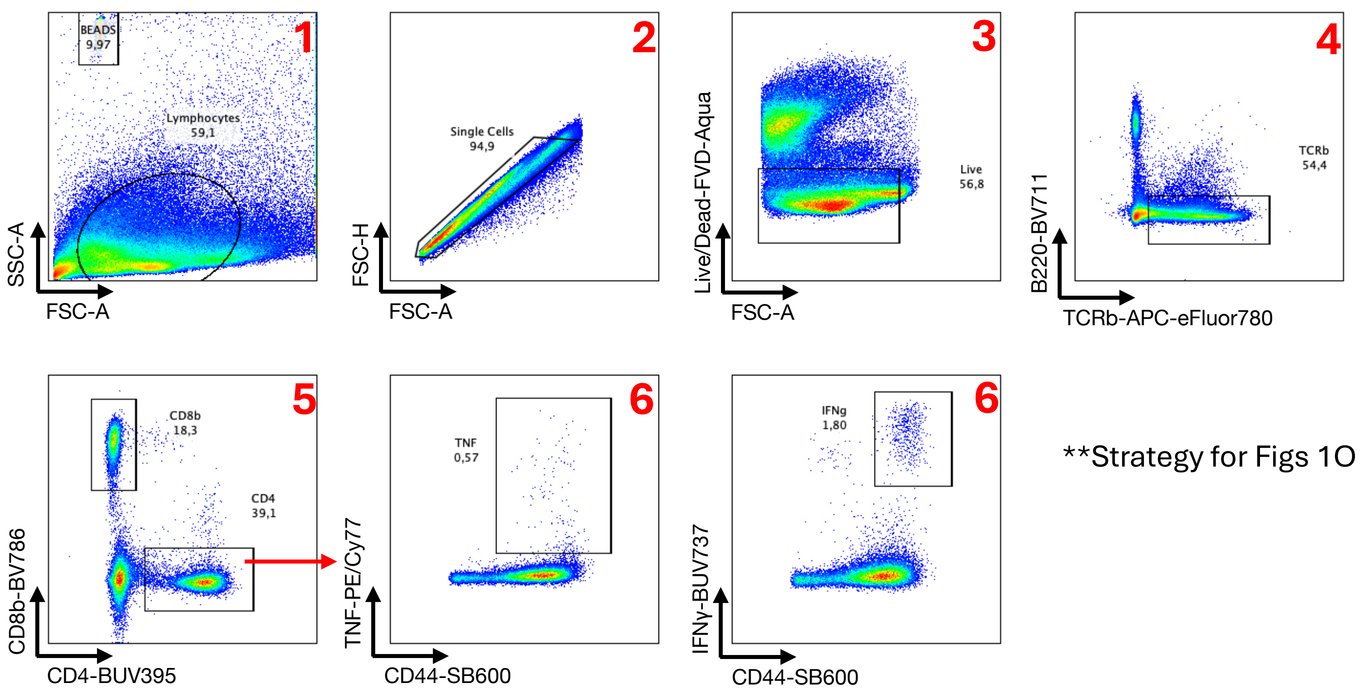

**Gating strategy for mLN and spleen B and T cells**

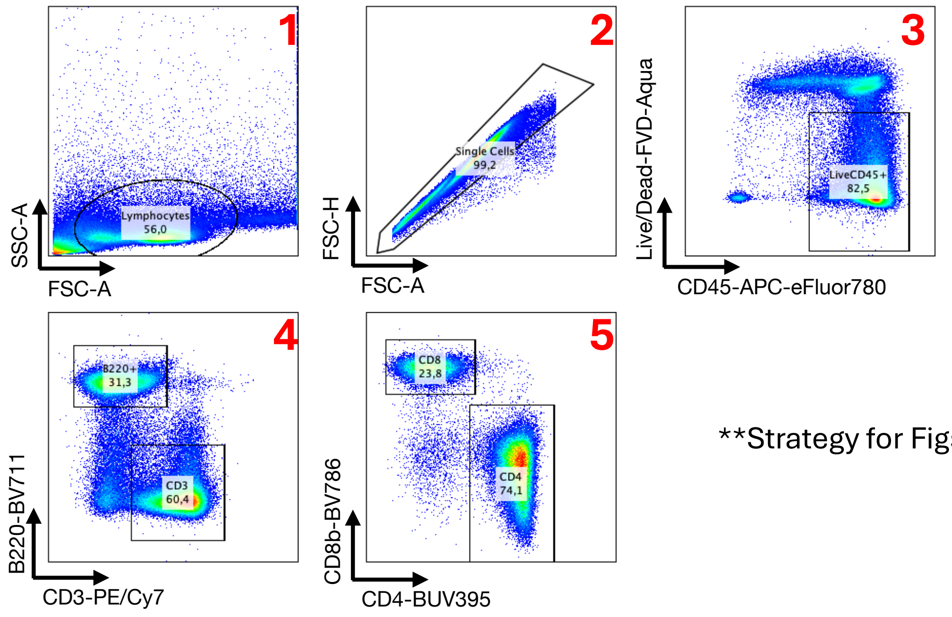


**Strategy for Figs 4B-E; S4B, S4D

**Gating strategy for mLN and spleen SMARTA T cells**

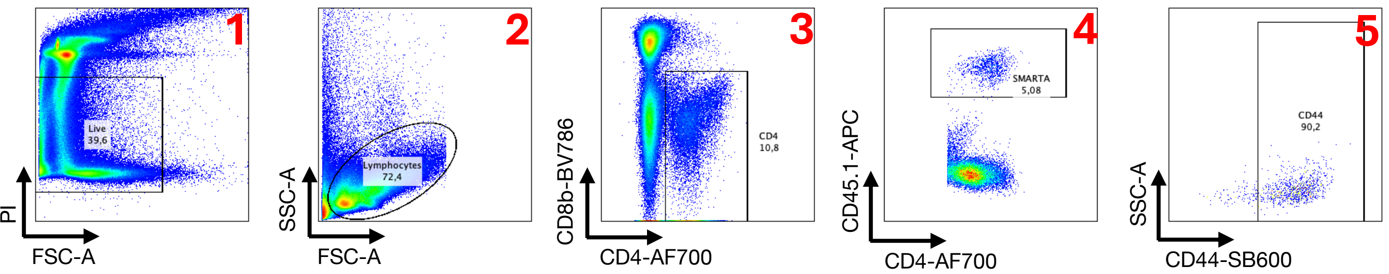


**Strategy for Figs 4G-M; S5A-G

#### Gating strategy for lamina propria stromal cells

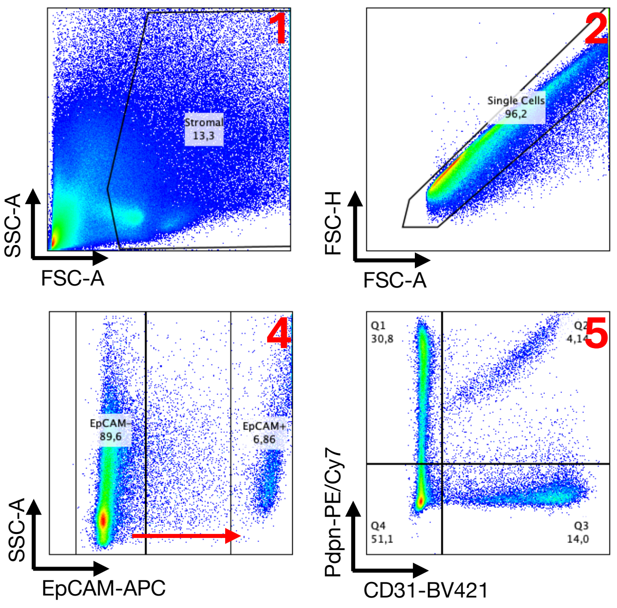


Q1 = Fib.
Q2 = LEC
Q3 = BEC
Q4 = DN

**Strategy for Figs 1K

#### Gating strategy for mLN stromal cells

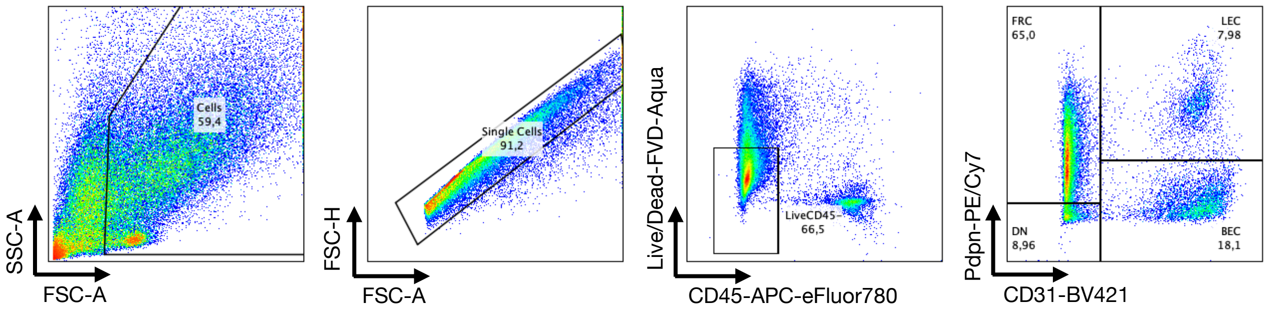


**Strategy for Figs 6N; S2A-E
